# The cryo-EM structure of the neurofibromin dimer reveals the molecular basis for von Recklinghausen disease

**DOI:** 10.1101/2021.02.18.431788

**Authors:** Christopher J. Lupton, Charles Bayly-Jones, Laura D’Andrea, Cheng Huang, Ralf B. Schittenhelm, Hari Venugopal, James C. Whisstock, Michelle L. Halls, Andrew M. Ellisdon

## Abstract

Neurofibromin (NF1) is a tumour suppressor mutated in neurofibromatosis type 1 (von Recklinghausen disease), one of the most common human genetic diseases(1). NF1 regulates cellular growth through suppressing the Rat Sarcoma (RAS) pathway and, accordingly, mutations in this protein drive numerous cancers, including melanoma, ovarian, breast and brain cancer(2, 3). Currently, however, the molecular basis for NF1 function remains to be understood. Here we address this problem and use cryogenic Electron Microscopy (cryo-EM) to determine the structure of fulllength NF1. The 640 kDa NF1 homodimer forms an extraordinary lemniscate (∞) shaped molecule that is ~30 nm in length and ~ 10 nm wide. Each NF1 monomer comprises an N-terminal HEAT-repeat domain (N-HEAT), a guanosine triphosphatase activating protein (GAP)-related domain (GRD), a Sec14 homologous and pleckstrin homologous module (SEC-PH), and a C-terminal HEAT domain (C-HEAT). The core NF1 scaffold is formed via a head-to-tail dimer of the N- and C-HEAT domains. This platform, which is responsible for interacting with more than 10 regulatory binding partners, comprises an extraordinary array of over 150 α-helices. Analysis of these EM data revealed that the GRD and SEC-PH domain are highly mobile with respect to the core scaffold and could not initially be accurately placed in electron density. Strikingly, however, using 3D variability analysis we were able to identify a significant subpopulation of NF1 particles and determine the complete NF1 structure to 5.6 Å resolution. These data revealed that the catalytic GRD and lipid binding SEC-PH domain are positioned against the core scaffold in a closed, autoinhibited conformation. We postulate that interaction with the plasma membrane may release the closed conformation in order to promote RAS inactivation. Our structural data further allow us to map the location of disease-associated NF1 variants and provide a long sought-after structural explanation for the extreme susceptibility of the molecule to loss-of-function mutations. Finally, it is suggested that approaches to combat NF1-linked diseases may include release of the autoinhibited state in order to improve NF1 catalytic efficiency.

---

The oncogenic RAS family of small GTP-binding proteins (Neuroblastoma RAS, Harvey RAS, and Kirsten RAS) have crucial roles in coordinating eukaryotic cell growth(4). Growth factor signalling promotes the cycling of RAS proteins from their inactive GDP-bound state to their active GTP-bound state. GTP-bound RAS proteins activate fundamental signalling pathways such as the RAF-MEK-ERK cascade and the PI3K-Akt-mTOR pathway(5). NF1 is a highly conserved RAS GAP that catalyses the hydrolysis of RAS-bound GTP(6). As such, NF1 acts as a potent off-switch for RAS-family signalling(2) (Extended Data Fig. 1).

Germline and sporadic mutations of *NF1* cause the common genetic condition neurofibromatosis type 1 (von Recklinghausen disease) with variable clinical presentations, including benign cutaneous and plexiform neurofibro-mas1. *NF1* is also a common driver gene in numerous cancers(7) with frequent mutations, deletions, or structural rearrangements in melanoma(8), breast cancer(9), lung cancer(10), and glioblastoma(11). Mutations that cause neurofibromatosis type 1 and cancer-associated somatic mutations are distributed throughout the *NF1* gene(12, 13) without widespread clustering to the catalytic GAP domain, suggesting an acute susceptibility of the entire protein to dysregulation. NF1 inactivation increases GTP-bound RAS levels driving downstream activation of MAPK and PI3K pathways and aberrant cell growth(14–16). NF1 loss further dysregulates many additional processes that lead to cognitive defects(17), conferral of drug resistance in cancer therapy(18), and learning defects(19).

NF1 interacts with a very large number of cellular binding proteins and effectors (Extended Data Fig. 1). Notably, NF1 binds Sprouty Related EVH1 Domain Containing 1 (SPRED1), an event that enables its membrane recruitment(20). The SEC-PH module is also postulated to play an additional role in membrane recruitment through lipid and phosphoinositide (PI) binding(21). Once at the membrane the catalytic NF1 GRD stabilises GTP-bound RAS to promote GTP-hydrolysis(22, 23). SEC-PH binding to G-protein βγcan also inhibit NF1, modulating RAS activation and neural pathways central to opioid addiction(24). Multiple further binding sites are distributed throughout NF1 including sites for tubulin binding in mitosis(25), 14-3-3 binding(26), as well as protein kinase A and C phospho-regulation(27).

To better understand NF1 function we expressed recombinant protein in insect cells and determined the cryo-EM structure of the full-length 640 kDa homodimer. The structure presented significant challenges with respect to particle heterogeneity, however, after symmetry expansion one wing of the NF1 homodimer could be reconstructed to a nominal resolution of 4.6 Å (Fig 1a, Extended Movie 1, Extended Data Fig. 2, 3, 4 and Extended Data Table 1). The maps were of sufficient quality to completely build and sequence de novo the N- and C-HEAT domains (Fig. 1b and Extended Data Fig. 5). In contrast, however, the central GRD and SEC-PH domains were poorly resolved and showed a high degree of mobility within the majority of particles. To address this, we used 3D variability analysis to identify a subpopulation of NF1 particles and determined the 5.6 Å structure of the complete NF1 molecule (Extended Data Fig. 4). In this latter reconstruction, clear secondary structure density was observed for the entire molecule (Fig. 1c-e) and crystal structures of both the GRD(20) and SEC-PH modules(21) could be fitted unequivocally into the density with each domain packed against the N- and C-HEAT domains. Extensive crosslinking mass spectrometry with BS^3^ and BS^2^G lysine crosslinkers confirm the placement of the central GRD and SEC-PH domains into the cryo-EM maps (Fig. 1f, g) as well as the sequence assignment of the dimeric N- and C-HEAT repeat core (Extended Data Fig. 5 and Extended Data Table 2). The GRD also crosslinks to known phospho-regulatory regions in NF1 (Extended Data Fig. 6).

**Fig. 1.**
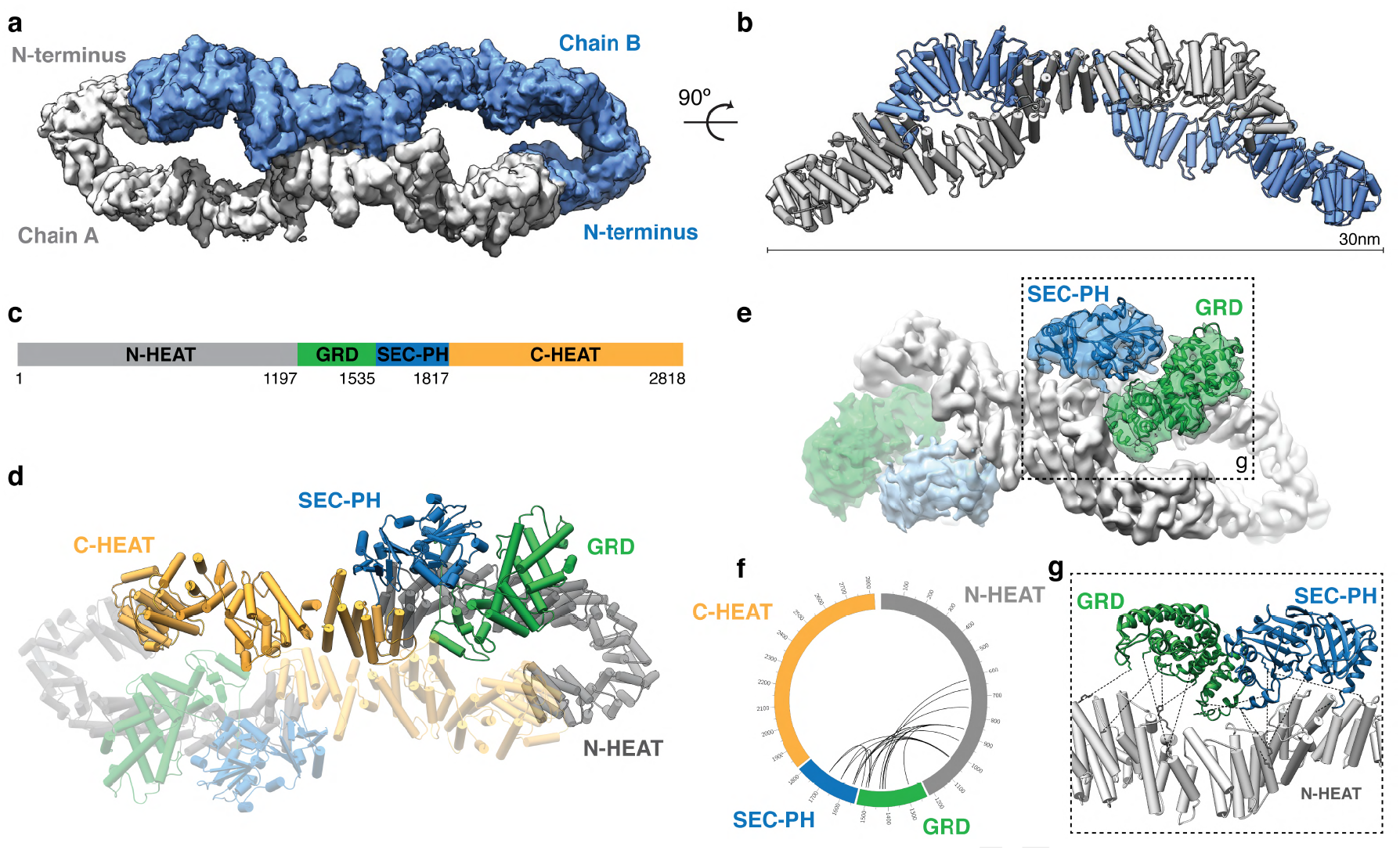
Structure of the NF1 homodimer. **a.** The 4.6 Å cryo-EM (composite) map of the NF1 core highlighting the intercalation of each monomer chain to form an extended lemniscate dimer. **b.** Overview of the NF1 dimer structure with **α**-helices as cylinders. **c.** Schematic of the NF1 domain layout. **d.** Overview of the domain organisation of the complete NF1 dimer structure with **α**-helices as cylinders. Overall, the N-HEAT domain forms a helicoidal arrangement of **α**-helices marked by the formation of a tight arch at each tip of the molecule. The N-terminal arch allows the N-HEAT domain to extend back towards the centre of the molecule. The GRD exits the central fold and runs back out along the edge of the N-HEAT domain before doubling back to form the SEC-PH module and re-joining the C-HEAT domain. The C-HEAT **α**-helical domain forms a central symmetrical dimer interface with the equivalent C-HEAT domain of the second dimer chain. The fold is closed at each end by two symmetrically equivalent interfaces that link the N- and C-termini of each dimer chain. **e.** The 5.6 Å cryo-EM map of the autoinhibited NF1 dimer oriented to display the fits of the atomic models of the GRD20 and SEC-PH21 domains. **f.** Intradomain BS^3^ and BS^2^G crosslinks detected between the N-HEAT, GRD, and SEC-PH domains (n=3). **g.** Boxed region from e shown to highlight the compatibility of crosslinks with the conformation of the GRD and SEC-PH domains.

Analysis of these structural data reveal that the NF1 scaffold comprises a large lemniscate-shaped assembly that forms as a consequence of head-to-tail dimerization of the N-and C-HEAT domains (Fig. 1a and Extended Movie 1). Each wing of the structure curves away from a central interface with an outer dimension spanning a remarkable ~30 nm between each tip (Fig. 1b). Despite being well resolved in the cryo-EM maps, the central scaffold is remarkably flexible with a large degree of continuous conformational heterogeneity detected across the dataset (Extended Movie 2). Docking sites of more than 10 known NF1 binding partners map across the entire NF1 structure (Extended Data Fig. 7). These data suggest, that like the karyopherin HEAT-repeat family(28), the core scaffold acts as a flexible interaction platform coordinating the many functions of NF1 that span from cell growth(2) to cell division(25).

NF1 dimer assembly is mediated by five interfaces. The central interface formed by the symmetrical and anti-parallel alignment of the C-HEAT domains, two symmetrically equivalent N- and C-terminal interfaces that close each end of the lemniscate HEAT-repeat fold, and two entry and exit interfaces that enable the flexible GRD and SEC-PH domains to align on the edge of the core scaffold (Fig. 2a, b). The central interface is extensive (~1670 Å^2^ buried surface area) and formed by multiple C-HEAT domain helices as well as the highly conserved αC2-3 loop that is sandwiched at each end of the symmetrical interface (Fig. 2a, c). The central interface is particularly well conserved and likely the main energetic driver of NF1 dimer assembly (Extended Data Fig. 8 and Extended Data Fig. 9).

**Fig. 2.**
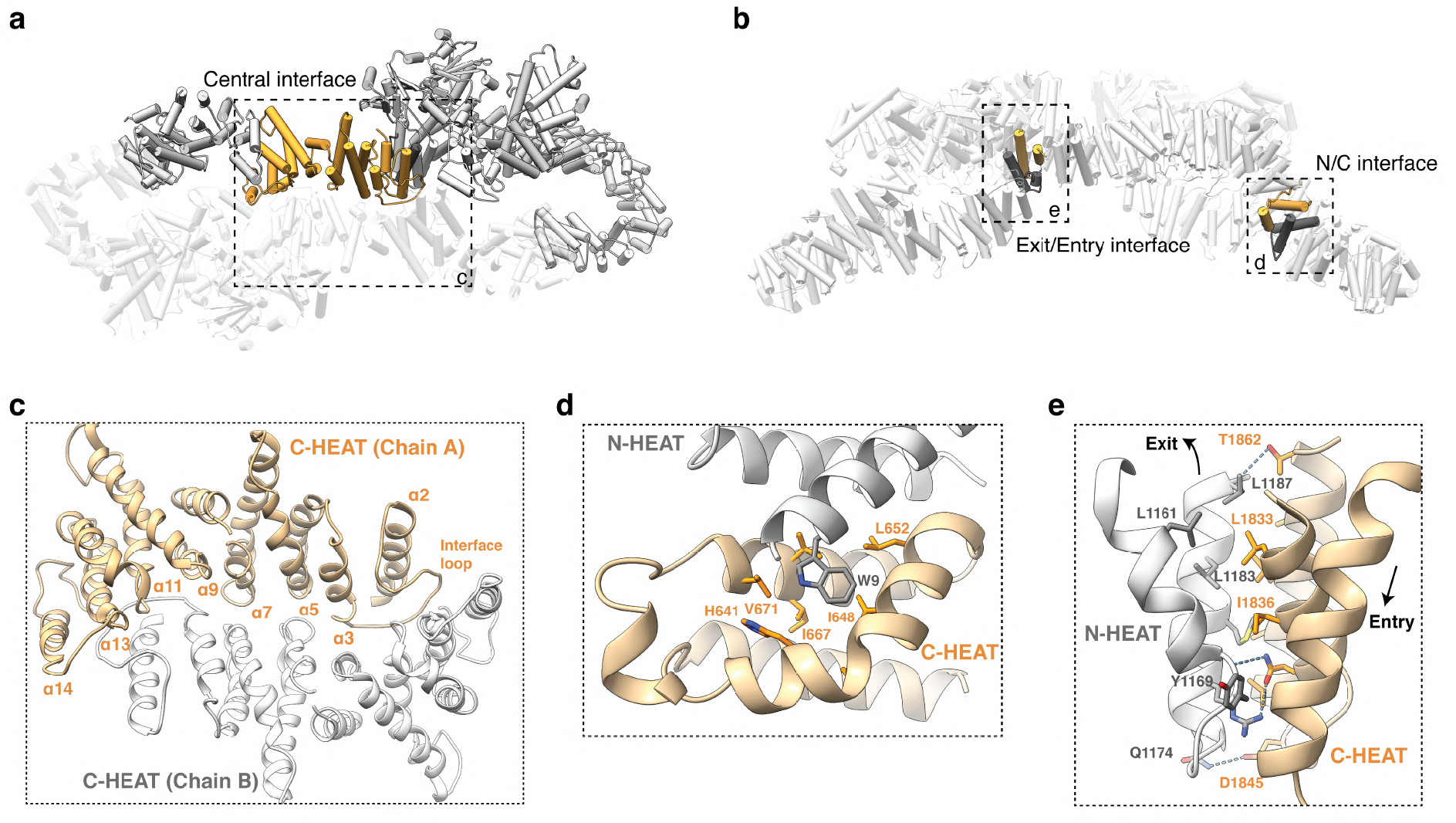
Structural basis of NF1 dimer assembly. **a.** The NF1 structure with the symmetrical central interface formed between the two C-HEAT domains highlighted. **b.** The NF1 structure with a single N/C interface and Exit/Entry interface highlighted. **c.** Boxed region from a with key interacting secondary structure elements indicated. **d.** Boxed region from b highlighting the W9 residue of the N-HEAT domain buried in the hydrophobic pocket of the C-HEAT domain. **e.** Boxed region of b highlighting key interacting residues in the continuous HEAT-repeat array interface of the N- and C-HEAT domains. Arrows indicate the direction of the exit and entry **α**-helices that support the placement of the GRD and SEC-PH domains on the core scaffold. Select amino acids are shown as sticks and indicated by single letter code. Potential hydrogen-bonds displayed as blue dotted lines.

The two smaller (~905 Å^2^ buried surface area) equivalent N- and C-HEAT interfaces are formed by the packing of helices αN1 and αN2 of the N-HEAT domain with helices αC32 and αC33 of the C-HEAT domain to form a continuous helicoid topology (Fig. 2d). The centre of the interface is predominantly hydrophobic, with the conserved N-HEAT tryptophan (W9) deeply buried into a hydrophobic pocket formed by helices αC32 and αC33 (Fig. 2d). The two entry and exit interfaces are formed by the N-HEAT helical pair αN44 and αN45 that form a continuous HEAT-repeat like array with the C-HEAT helical pair αC1 and αC2. A total of ~850 Å^2^ is buried in each interface with predominantly hydrophobic packing between helices (Fig. 2e). This interface is central to supporting the integrity of the central scaffold as well as the flexible positioning of the GRD and SEC-PH domains on the scaffold edge.

In the 5.7 Å structure of the complete NF1 molecule, the GRD and SEC-PH modules pack tightly against the N- and C-HE AT domains. Whilst resolution limited detailed analysis, the GRD sits buried in the cleft formed by the N- and C-HEAT domains making contacts with both conserved C-HEAT domain α-helices as well as surface loops of the N-HEAT repeats (Fig. 3a-c). Analysis of the closed NF1 structure reveals that the RAS-binding interface is oriented towards the central scaffold in a conformation that is sterically incompatible with direct RAS binding (Fig. 3a, b). Similarly, the lipid binding sites of both the SEC and PH domains(21) are also occluded and would need to undertake significant rotation to enable membrane binding (Fig. 3d, e). In contrast, the modelled SPRED1 binds intimately to NF1 surface potentially interfacing with a previously unrecognized site in the C-HEAT domain (Fig. 3b, c) providing a clear route for SPRED1-driven membrane recruitment(29).

**Fig. 3.**
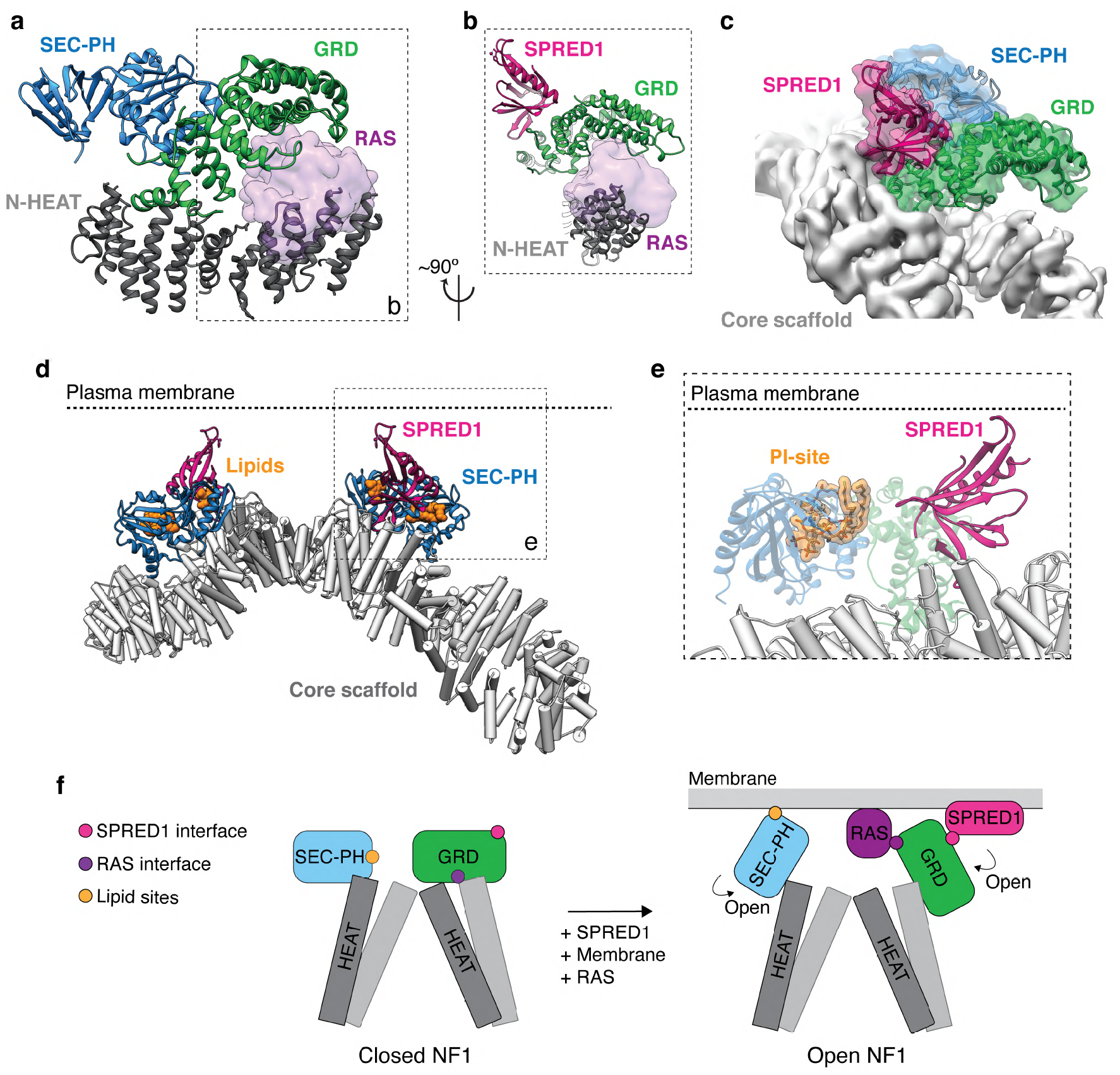
The GRD and SEC-PH domains are in a closed conformation. **a.** RAS binding to the GRD is sterically occluded by the N-HEAT domain. **b.** Boxed region from **a** rotated to highlight steric clashes of RAS with the N-HEAT domain and the SPRED1 binding site exposed on the GRD domain. **c.** NF1 oriented to highlight the close association of the modelled SPRED1 domain to the central scaffold. **d.** The NF1 structure oriented to highlight the horizontal alignment of the SEC-PH lipid binding pockets as well as the SPRED1 binding region on each GRD. The predicted plasma membrane position is indicated to highlight the symmetrical alignment of known NF1 membrane binding regions. **e.** Boxed region from d highlighting the buried PI binding site in the PH domain. **f.** Proposed regulatory model of NF1. A subpopulation of NF1 is in a stable closed conformation marked by the steric occlusion of the GRD interface and lipid-binding sites. The SPRED1 binding site(20) is exposed on the surface of the closed GRD conformation and is known to recruit NF1 to the membrane(29). Conformational rearrangements of the GRD and SEC-PH modules are required for RAS and lipid binding, respectively. For clarity, the model displays only a GRD or SEC-PH domain on each side of the core scaffold.

Collectively these data suggest that we have identified a closed conformation of NF1. Our data further suggest that significant rearrangement of both the GRD and SEC-PH domains must take place upon SPRED1 membrane recruitment in order to enable RAS turnover and lipid binding, respectively (Fig. 3e). Consistent with this idea, our structural data reveal that GRD and SEC-PH domains are highly mobile and can thus readily transition between the closed conformation and an open conformation where the GRD and SEC-PH domains are exposed and flexible.

Lastly, in order to gain understanding of the molecular basis of neurofibromatosis type 1 we analysed the frequency of disease-associated mutations across the *NF1* gene(13, 30) (Fig. 4a.). Neurofibromatosis type 1 mutations are evenly spread throughout the gene with only sporadic clustering evident along its length. Analysis of the COSMIC(12) database also highlights that somatic cancer-associated mutations are evenly distributed throughout the *NF1* gene (Fig. 4b). We mapped these disease-associated mutations onto the NF1 structure to gain greater insight into the mechanistic basis of loss-of-function of this key tumour suppressor protein (Fig. 4c). For example, the clinically significant neurofibromatosis type 1 mutants (ΔM991(31)) and (ΔL844-G848(32)) map to the N-HEAT domain and would likely interfere with helical packing directly beneath the SEC-PH interface (Fig. 4d). In addition, the common cancer-associated mutation R1849Q(33) packs intricately against the central interface (αC2-3) loop, a crucial region for NF1 dimer assembly (Fig. 4e). However, globally, NF1 mutations do not cluster to any one interface or region of the protein but are uniformly distributed throughout the structure (Fig. 4c). It is clear from the intricate NF1 fold that missense, deletion, or nonsense mutations are likely to have an acute effect on the stability of the entire NF1 protein fold. For example, any mutation that disrupts the intricate HEAT-repeat packing of the core scaffold may result in an inability of the lemniscate-fold to close and downstream loss of function.

**Fig. 4.**
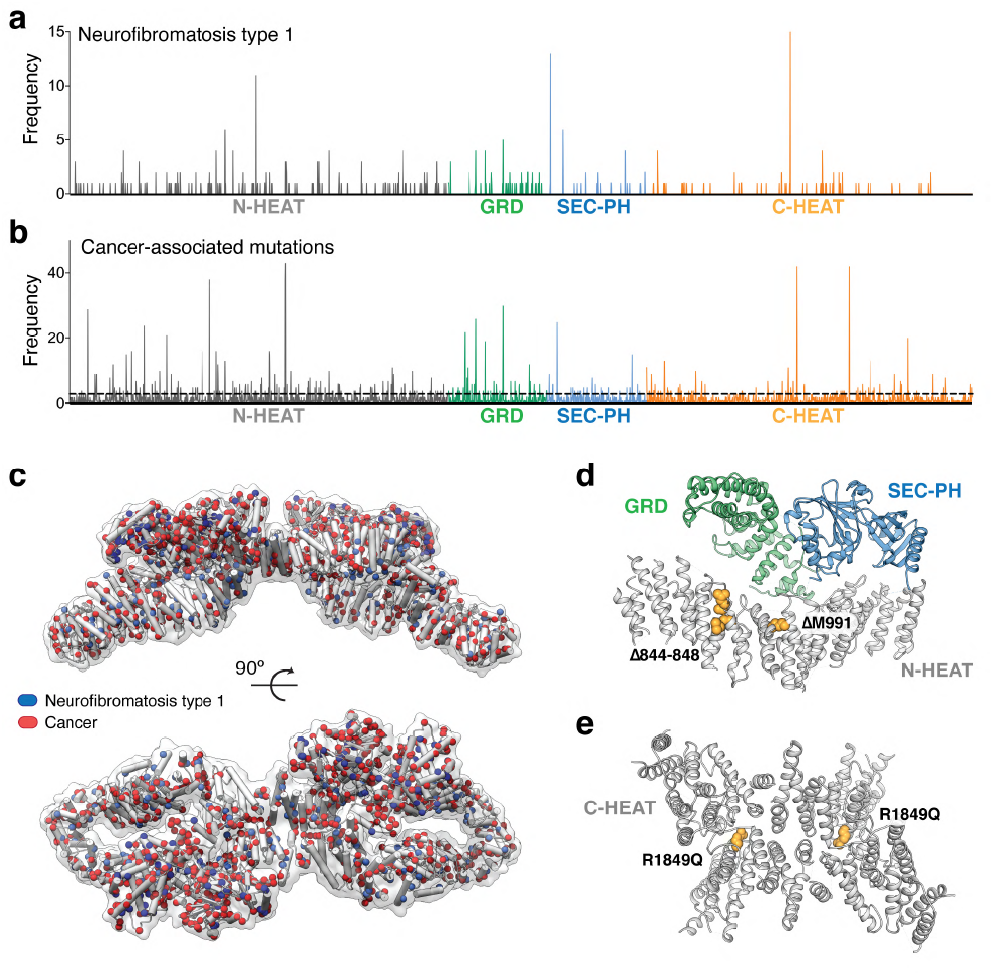
Distribution of neurofibromatosis type 1 and cancer-associated NF1 mutations. **a.** Frequency of neurofibromatosis type 1 (von Recklinghausen disease) mutations from two patient studies(13, 30). **b.** Frequency of NF1 cancer-associated mutations found in the COSMIC(12) database mapped across the NF1 gene. Sites of missense, nonsense, and deletions are included in the analysis to highlight the lack of clustering. Positioning of the NF1 domains from the cryo-EM structure are indicated **c.** The NF1 structure with C**α** atoms of residues mutated in neurofibromatosis type 1 (blue) or cancer (red) displayed as spheres. All neurofibromatosis type 1 mutations from the indicated studies are displayed and for clarity cancer-associated mutations are only displayed if they were in the COSMIC12 database three or more times. **d.** Indicate neurofibromatosis type 1 mutants are within the N-HEAT domain close to the GRD interface. **e.** Location of the cancer-associated R1849Q mutant within the central NF1 dimer interface.

This sensitivity of the NF1 fold to mutations also rationalises the lack of mutational hot spots within the tumour suppressor.

In conclusion, we have shown that NF1 assembles into a large homodimer characterised by a central lemniscate-shaped core formed by the assembly of the N- and C-HEAT domains. NF1 dimer assembly requires the coordinated packing of eight domains, which contain over 150 α-helices, to form the final topology. This highly mobile molecular scaffold provides an elongated binding platform for both the GRD and SEC-PH domains as well as the many NF1 cellular effector molecules that have been identified to date. We have further identified a sub-population of NF1 with the catalytic and lipid binding domains in a closed conformation suggesting a previously unidentified mechanism of NF1 autoinhibition. This pivotal role of the core scaffold in coordinating both NF1 activity and regulation provides a rationale for the evolutionary conservation of this unusually complex fold.

Finally, our work provides a long-sought after molecular and structural context for NF1 dysregulation in disease. As such, mutation or deletion at a disproportionate number of sites is likely to cause mis-assembly of the dimer and acute loss-of-function. We propose that the complex nature of the NF1 fold is thus a major contributing factor for the acute sensitivity of the *NF1* gene to mutations in disease. However, given the identification of the closed conformation of NF1 we hypothesise that developing molecules to stabilize the open conformation may represent a strategy to rescue NF1 loss-of-function in disease where an active allele remains.

## Material and methods

### Protein expression and purification

NF1 (isoform 1) was subcloned from R777-E139 Hs.NF1 (a gift from Dominic Esposito, Addgene plasmid #70423) as an N-terminal His_6_-tagged protein in pFastBac1 (Invitrogen). NF1 was expressed in Sf9 cells for 2 days following baculovirus infection (Bac-to-Bac, Invitrogen). Cells were harvested by centrifugation and lysed by sonication in 20 mM Tris-HCl pH 8.0, 500 mM NaCl, 10 % (v/v) glycerol, and 5 mM β-mercaptoethanol. Lysates were cleared by centrifugation, filtered at 0.45 μm, and incubated with Ni-NTA resin (Qiagen) and 20 mM imidazole at 4 °C with agitation. The resin was washed with lysis buffer, and NF1 eluted with 500 mM imidazole in the same buffer. NF1-containing fractions were further purified on a Superose 6 10/300 GL size-exclusion column (Cytiva) equilibrated in 20 mM HEPES pH 8.0, 150 mM NaCl, and 2 mM DTT.

Site-directed mutagenesis was performed on KRAS 1-169 (Q61H, isoform b, a gift from Cheryl Arrowsmith, Addgene plasmid #25153) to obtain the wild-type sequence. N-terminally His6-tagged KRAS was expressed in BL21-CodonPlus(DE3)-RIL cells (Stratagene) by isopropyl-β-D-1-thiogalactopyranoside induction at 18 °C. Cells were lysed by sonication in 20 mM Tris-HCl pH 8.0, 500 mM NaCl, 10 % (v/v) glycerol, and 3 mM β-mercaptoethanol, with two complete EDTA-free protease inhibitor tablets (Roche). Cells were clarified by centrifugation, filtered through a 0.45 μm membrane, and incubated with Ni-NTA resin (Qiagen) and 20 mM imidazole at 4 °C with agitation. The resin was washed with lysis buffer and KRAS was eluted in lysis buffer containing 300 mM imidazole. KRAS-containing fractions were further purified on a HiLoad Superdex 75 16/60 (Cytiva) pre-equilibrated with 20 mM HEPES pH 8.0, 150 mM NaCl, 10 % (v/v) glycerol, and 2 mM DTT.

### Cryo-EM sample preparation and data collection

Quantifoil R1.2/1.3 200 mesh copper grids were glow discharged for 90 s at 15 mA using a Pelco easiGlow instrument. Freshly purified NF1 (3 μL at 0.5 mg ml^−1^) was applied to the freshly discharged grid and vitrified in liquid ethane using a Vitrobot Mk IV (Thermo Fisher Scientific) with a blot of force of −2 for 2.5 s. Temperature and relative humidity were maintained at 4 °C and 100 %, respectively. Data were collected on a Talos Arctica (Thermo Scientific) operating at 200 kV with a 50 μm C2 aperture. Micrographs were acquired using a bottom mounted Falcon 3 direct electron detector in counting mode at a nominal magnification of 120,000×, corresponding to a calibrated physical pixel size of 1.194 Å. The electron dose rate was set to 0.8 electrons pixel^−1^ s^−1^ with a total exposure time of 71.68 s, yielding a total dose of 40 electrons Å^−2^. Automated collection was carried out using EPU with beam-shift to collect 9 images per stage movement. Defocus range was set between −0.5 to −2.2 μm.

### Cryo-EM data processing

All initial data monitoring and micrograph denoising were performed in Warp(34) (v1.0.9) to assess grid quality and select appropriate regions for data collection. An initial pass of the data was performed in cryoSPARC(35) (v2.14.2). Here, particle picking was performed by the elliptical blob method, these 23,480 particles were extracted in a box of 432 pixels, down sampled to 2.388 Å pixel^−1^ and subjected to 2D classification. An *ab initio* model was generated from 1,818 particles without imposing symmetry.

Upon completion of data collection, 2,675 dose-fractionated movies were corrected for beam induced motion and compensated for radiation damage within MotionCor2(36) (v1.1.0). All aligned movie frames were subsequently summed into dose-weighted averages for further processing. The contrast transfer function parameters were re-estimated with CTFFIND(37) (v4.1.8). Curated particles from the initial pass were used to train a crYOLO(38) (v1.6.1) model which was used to pick 192,225 particles from the full dataset. These were subjected to multiple rounds of 2D classification in both RELION(39) (v3.1) and cryoSPARC, yielding 95,564 particles of sufficient quality and homogeneity. A consensus reconstruction was generated by refining this subset in RELION with C2 symmetry yielding an 8.2 Å map after polishing and re-sampling to 1.194 Å pixel^−1^. These particles were 3D classified in RELION without further alignments. A single subset was refined using RELION and SIDESPLITTER(40) yielding a map of 7.5 Å resolution. Further local refinement in cryoSPARC lead to a final dimer reconstruction at 6.2 Å resolution.

CryoSPARC 3D variational analysis and RELION multibody analysis was performed on the consensus reconstruction to assess the degree of flexibility and guide signal subtraction. These analyses revealed significant movement of the two NF1 wings, as well as flexibility within individual lobes. Therefore, symmetry expansion of the consensus reconstruction was performed, and each wing was re-extracted after signal subtraction of the other wing. This gave rise to 191,128 sub-particles in a 256-pixel box. An additional round of 2D classification was performed to remove empty and junk particles. Refinement in RELION yielded a 6.4 Å map after symmetry expansion and signal subtraction, indicating flexibility of the NF1 homodimer had significantly limited the resolution of the consensus reconstruction. Further 3D classification of these particles yielded two homogenous subsets. Refinement of the CTF parameters was performed in cisTEM(41) with a high-resolution cut-off maintained above the fall-off of the FSC. Lastly, a final homogenous refinement was performed with RELION and SIDESPLITTER yielding a 4.56 Å resolution map. Conversions between software were performed with EMAN(42) (v2.2), with code written in-house or by pyem. The FSC was used to estimate resolution at the 0.143 threshold. Local resolution was estimated by the windowed blocres FSC method as implemented in cryoSPARC. Map sharpening was performed in RELION, an ad hoc B-factor of −80 Å^−2^ was selected by inspection of the map.

A second round of 3D variation analysis was performed on the symmetry expanded particles, which after 3D principal component analysis, revealed a small discrete cluster of particles corresponding to an autoinhibited state of NF1. This subpopulation of 9,283 particles was re-extracted at the original pixel size and locally refined with gaussian priors to prevent diverging from previous alignment parameters. This analysis resulted in a 5.6 Å resolution map of the autoinhibited NF1 with GRD and SEC-PH domains with clearly resolved secondary structure.

The model of the N-HEAT and C-HEAT domains was built manually into the cryo-EM map using Coot(43), with iterative cycles of refinement carried out using PHENIX(44) and Isolde(45). Secondary structure and adaptive network constraints were employed until the final round of refinement, where only secondary structure constraints were imposed. The structure had excellent final geometry (Extended Data Table 1). Structures of both the GRD(20) (PDB: 6V65) and SEC-PH(21) (PDB: 3PG7) domains were rigid body fit into the 5.7 Å cryo-EM reconstruction with excellent agreement between the EM and crystal structures to produce an autoinhibited NF1 model. A final, full NF1 dimer model was generated by rigid body fitting the autoinhibited NF1 model into the 6.2 Å dimer reconstruction.

### CLMS

Crosslinking mass spectrometry was performed by adding either BS^3^ (Thermo Fisher) or BS^2^G (Thermo Fisher) crosslinkers at a 1:100 molar ratio to 4 μM NF1 in 20 mM HEPES pH 8.5, 150 mM NaCl, and 2 mM DTT. For BS^3^, samples were incubated at room temperature for 20 min prior to the addition of 50 mM Tris-HCl pH 8.0 to quench the reaction. For BS^2^G, samples were incubated at room temperature for 30 min prior to the addition of 20 mM NH_4_HCO_3_ to quench the reaction. For both crosslinkers, samples were snap-frozen in liquid nitrogen prior to further processing. Samples were subsequently denatured for 30 min at 65 °C in the presence of 10 mM DTT. Subsequently, 40 mM chloroacetamide was then added to the samples prior to incubation for 20 min at room temperature in the dark. A 1:100 (w/w) ratio of trypsin was added to the samples and further incubated at 37 °C overnight. Digestion was stopped with the addition of 1 % (v/v) formic acid. Samples were subsequently cleaned using OMIX C18 pipette tips (Agilent Technologies) and stored in 0.1 % (v/v) formic acid prior to mass spectrometry.

Samples were analysed by LC-MS/MS using a Dionex Ultimate 3000 RSLCnano system coupled onto an Orbitrap Fusion Tribrid instrument (Thermo Fisher). An Acclaim PepMap RSLC analytical column (75 μm × 50 cm, nanoViper, C18, 2 μm, 100 Å; Thermo Scientific) and an Acclaim PepMap 100 trap column (100 μm × 2 cm, nanoViper, C18, 5 μm, 100 Å; Thermo Scientific) were used to separate tryptic peptides by increasing concentrations of 80 % (v/v) acetonitrile (can) / 0.1 % (v/v) formic acid at a flow of 250 nl min^−1^ for 90 min. The mass spectrometer was operated in data-dependent mode with the following parameters. The cycle time was controlled for 3 s. The MS1 resolution was set at 120,000 and scan range of 375-2000 m/z. The AGC target was set at 1.0×10^6^ with an injection time of 118 ms. The MS2 resolution was set at 60,000 and the AGC target was set at 4.0×10^5^ with an injection time of 118 ms. pLink and pLink2 were used to identify BS^3^- or BS^2^G-crosslinked peptides(46). Each CLMS dataset is derived from three repeats and crosslinked peptides were analysed if they were identified at least twice with a P value of less than 10^−4^ and were greater than 10 residues apart. Visual representations of crosslinked peptides were generated in Circos(47).

#### GAP-activity assays

Assays for GAP-stimulated GTP hydrolysis were performed using the GTPase-Glo kit (Promega) based on the protocol by Mondal et al(48). Briefly, KRAS (wild-type) was added to white opaque 384-well plates (Corning) at a final concentration of 0.5 μM and GTPase reactions were initiated by adding NF1 at a final concentration of 0.5 μM (final volume 40 μL with 5 μM GTP added to buffer). Samples were incubated for 90 min at room temperature and luminescence measured using a BMG CLARIOstar plate reader with emission at 545-550 nm. In each case, we performed three independent experiments in duplicate. Data are expressed as the mean ±standard error of the mean.

#### Sedimentation velocity analytical ultracentrifugation

Sedimentation velocity analytical ultracentrifugation experiments were performed with full-length NF1 at 1 μM in 20 mM HEPES pH 8.0, 150 mM NaCl and 1 mM DTT on a Beckman Coulter Optima analytical ultracentrifuge with An60-Ti rotor at 25,000 rpm at 21 °C. Absorbance data was collected at 280 nm, and all data and frictional ratio calculations were analysed in SEDFIT(49). Buffer density, buffer viscosity and sample partial specific volumes were calculated based on their composition in SEDNTERP.

## DATA AVAILABILITY

The three-dimensional cryo-EM density maps are deposited into the Electron Microscopy Data Bank (https://www.ebi.ac.uk/pdbe/emdb/) under accession number XXX. The coordinates are deposited in the Protein Data Bank (https://www.rcsb.org) with accession number XXX. Mass spectrometry proteomics data have been deposited to the ProteomeXchange Consortium via the PRIDE(50) partner repository with the dataset identifier PXD023593.

## ACKNOWLEDGEMENTS

M.L.H. is a Viertel Senior Medical Research Fellow supported by The Cross Family and The Frank Alexander Charitable Trusts. J.C.W. is an Australian Research Council Laureate Fellow and honorary National Health and Medical Research Council of Australia Senior Principal Research Fellow. This research was supported by a NHMRC Project Grant to M.L.H. (APP1121029) and equipment funded by an Australian Research Council Grant (LE170100016). CBJ acknowledges the support of the Australian Government by way of an RTP stipend. The authors acknowledge the use of instruments and assistance at the Monash Ramaciotti Centre for Cryo-Electron Microscopy, a Node of Microscopy Australia. We also acknowledge the office of the Vice-Provost for Research and Research Infrastructure (VPRRI) at Monash University and of Bioplatforms Australia (BPA) as part of the National Collaborative Research Infrastructure Strategy (NCRIS).This preprint was formatted in LATEX using an adaptation of Ricardo Henriques’ template.

## AUTHOR CONTRIBUTIONS

A.M.E. and M.L.H. conceived the study. C.J.L. and L.D. performed cloning, protein expression, and purification. C.J.L. prepared cryo-EM grids. C.J.L. and H.V collected cryo-EM data. C.B-J., C.J.L. and A.M.E. processed cryo-EM data, built, and refined atomic models. C.J.L. and L.D. performed GAP activity assays. L.D., C.H. and R.B.S. performed crosslinking and mass spectrometry. All authors contributed to analyses of data and writing of the manuscript.

---

**Fig. S1.**
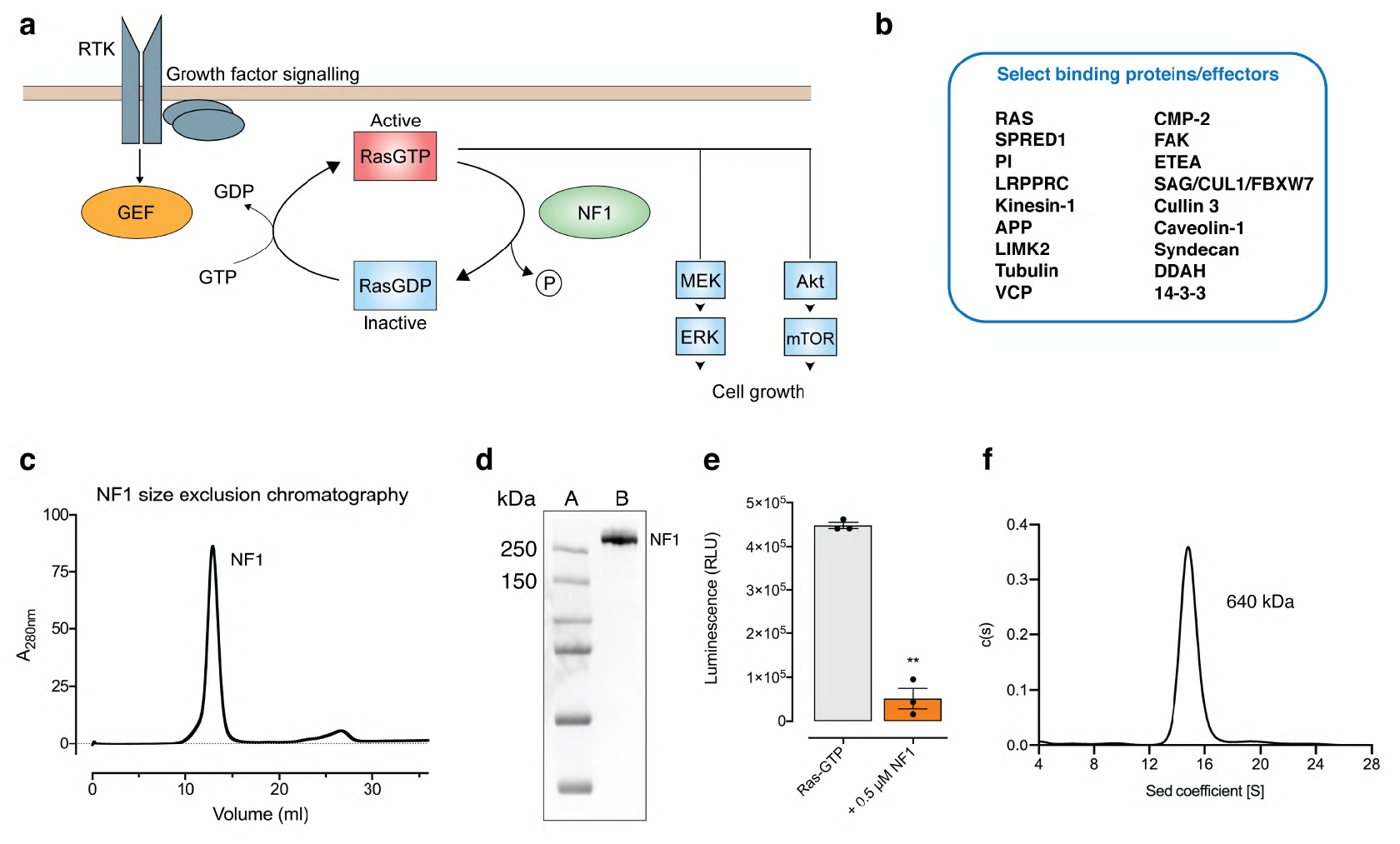
Purification and biochemical characterisation of the NF1 dimer. **a.** Schematic of the inhibitory role of NF1 in RAS signalling. **b.** Select NF1 binding partners and cellular effectors are listed(2). **c.** NF1 purifies as a single peak by size-exclusion chromatography. **d.** Purified recombinant NF1 is highly pure by SDS-PAGE. **e.** NF1 accelerates the hydrolysis of GTP by KRAS. Hydrolysis was measured using a GTPase-Glo kit (Promega) to detect the amount of GTP remaining in the reaction after incubation. ** p<0.01 versus Ras-GTP alone, paired two-tailed t-test. **f.** Sedimentation velocity analytical ultracentrifugation analysis indicates that NF1 likely forms a homodimer in solution (calculated molecular weight of 640 kDa, r.m.s.d. 0.004).

**Fig. S2.**
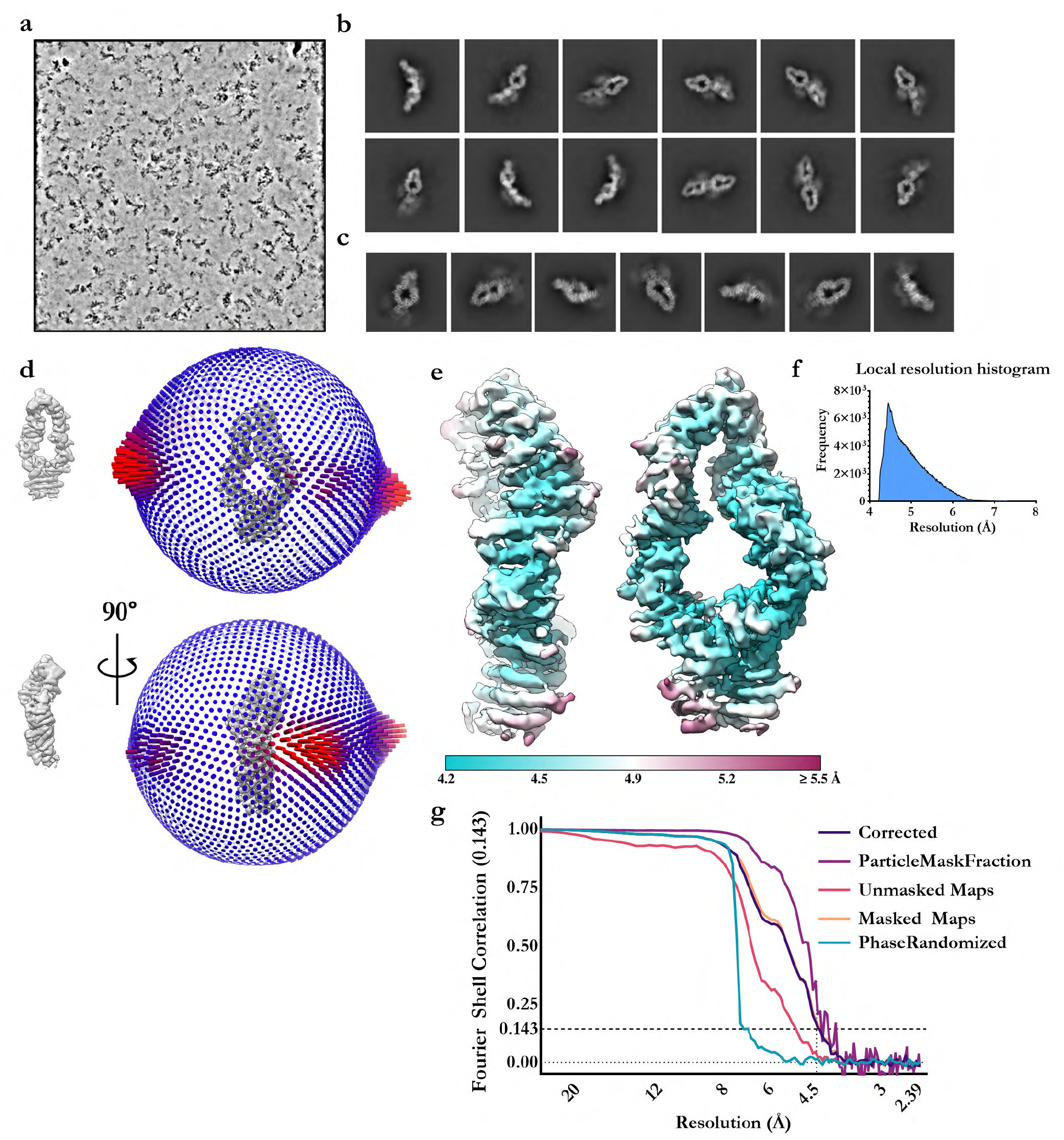
Cryo-EM summary and validation for the NF1 wing localised reconstruction. **a.** A representative denoised micrograph of NF1 collected on a Talos Arctica 200 kV **b.** Selected 2D class averages of the NF1 dimer. Diffuse signal can be observed at the central interface corresponding to flexible GRD/SEC/PH domains. **c.** Select 2D class averages of symmetry expanded, localised reconstructions of a single NF1 dimer lobe. **d.** Angular distribution of the final localised reconstruction, shown beside is the corresponding orientation of the reconstruction for clarity. **e.** The NF1 dimer lobe coloured by local resolution. **f.** Distribution of voxels as a function of local resolution. The majority of voxels are resolved to a mode value of ~4.5 Å. **g.** The Fourier shell correlation plot estimates a global resolution of 4.5 Å.

**Fig. S3.**
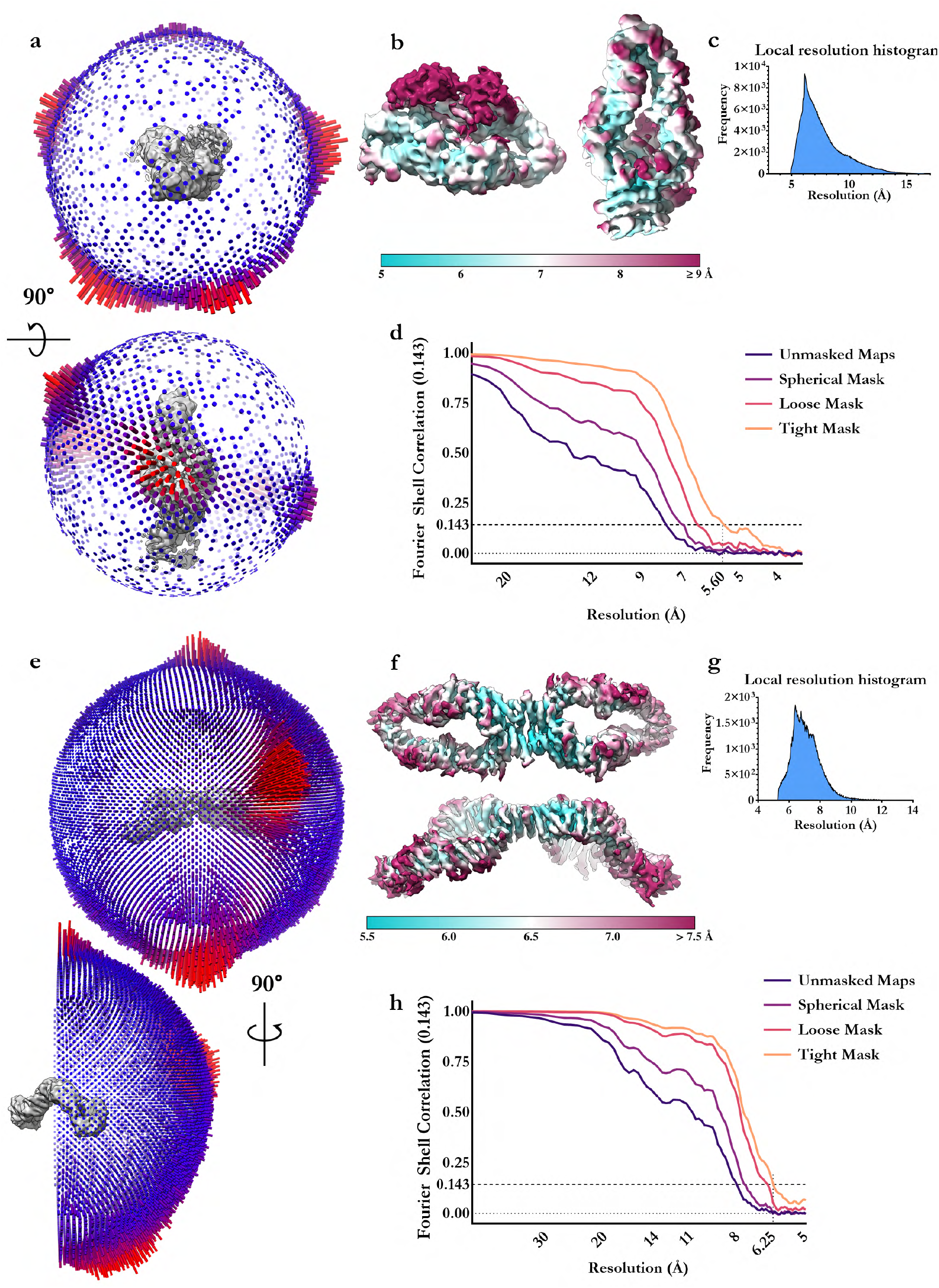
Cryo-EM summary and validation for the reconstructions of NF1 autoinhibited and dimeric states. **a.** Angular distribution of the autoinhibited NF1 localised reconstruction, **b.**, and local resolution analysis. **c.** Frequency distribution of voxels as a function of local resolution for the autoinhibited NF1 localised reconstruction. **d.** The Fourier shell correlation plot estimates a global resolution of 5.6 Å. **e, f, g.** As in **a**, **b**, **c**. for the full NF1 homodimer. **h.** The Fourier shell correlation plot estimates a global resolution of 6.3 Å.

**Fig. S4.**
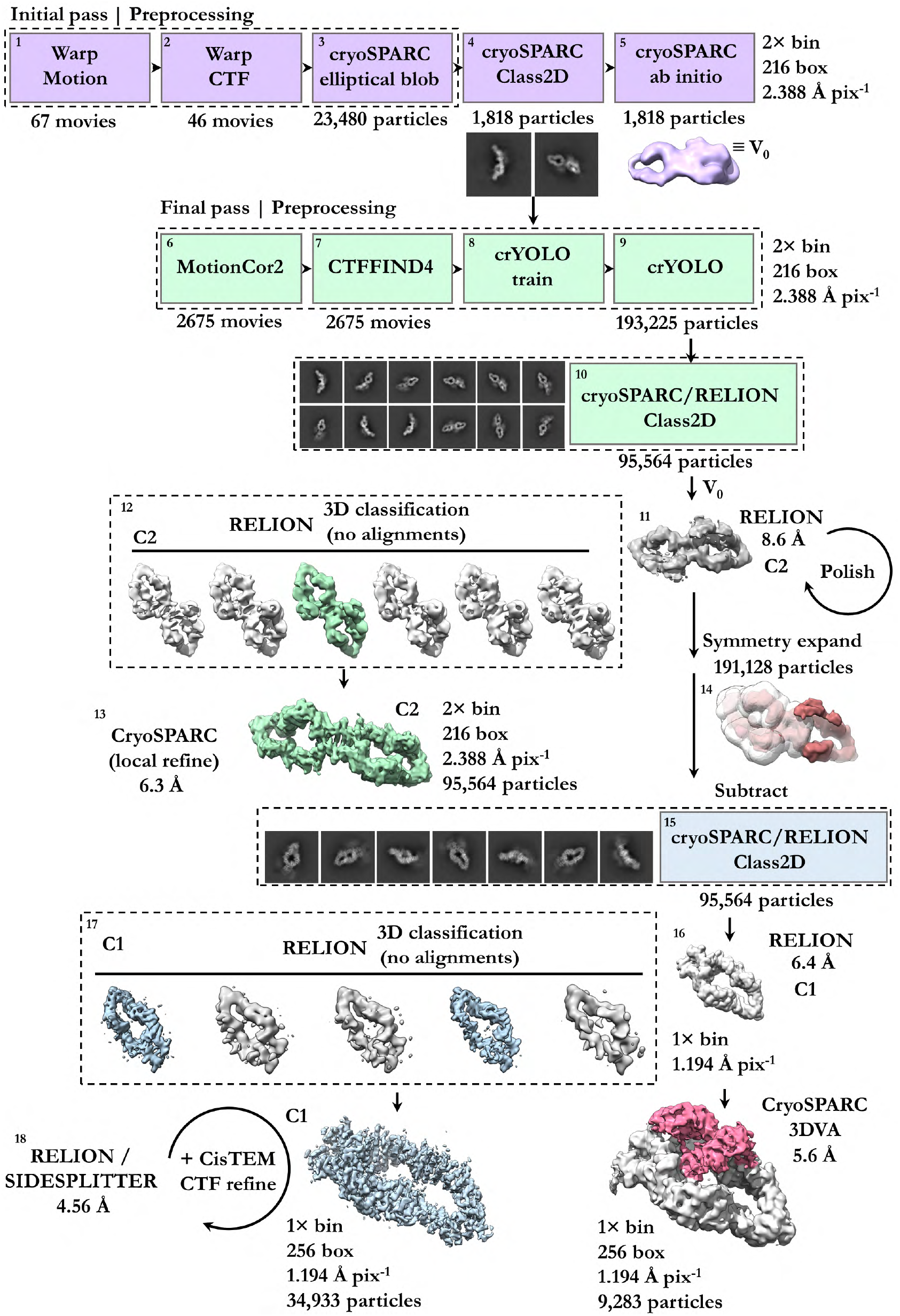
Cryo-EM data analysis and flowchart. Initial cryo-EM analysis is indicated in purple. Subsequent analysis of the whole dataset give rise to a 7.5 Å reconstruction of the NF1 dimer (green). Symmetry expansion and signal subtraction were utilised to generate a focused refinement of a single NF1 lobe (light blue) to overcome significant continuous conformational heterogeneity. The final 4.6 Å reconstruction was later fitted into the NF1 dimer reconstruction to yield a composite map. Variational analysis revealed a small discrete population of NF1 particles with resolved GRD and SEC-PH domains (red) in an autoinhibited conformation.

**Fig. S5.**
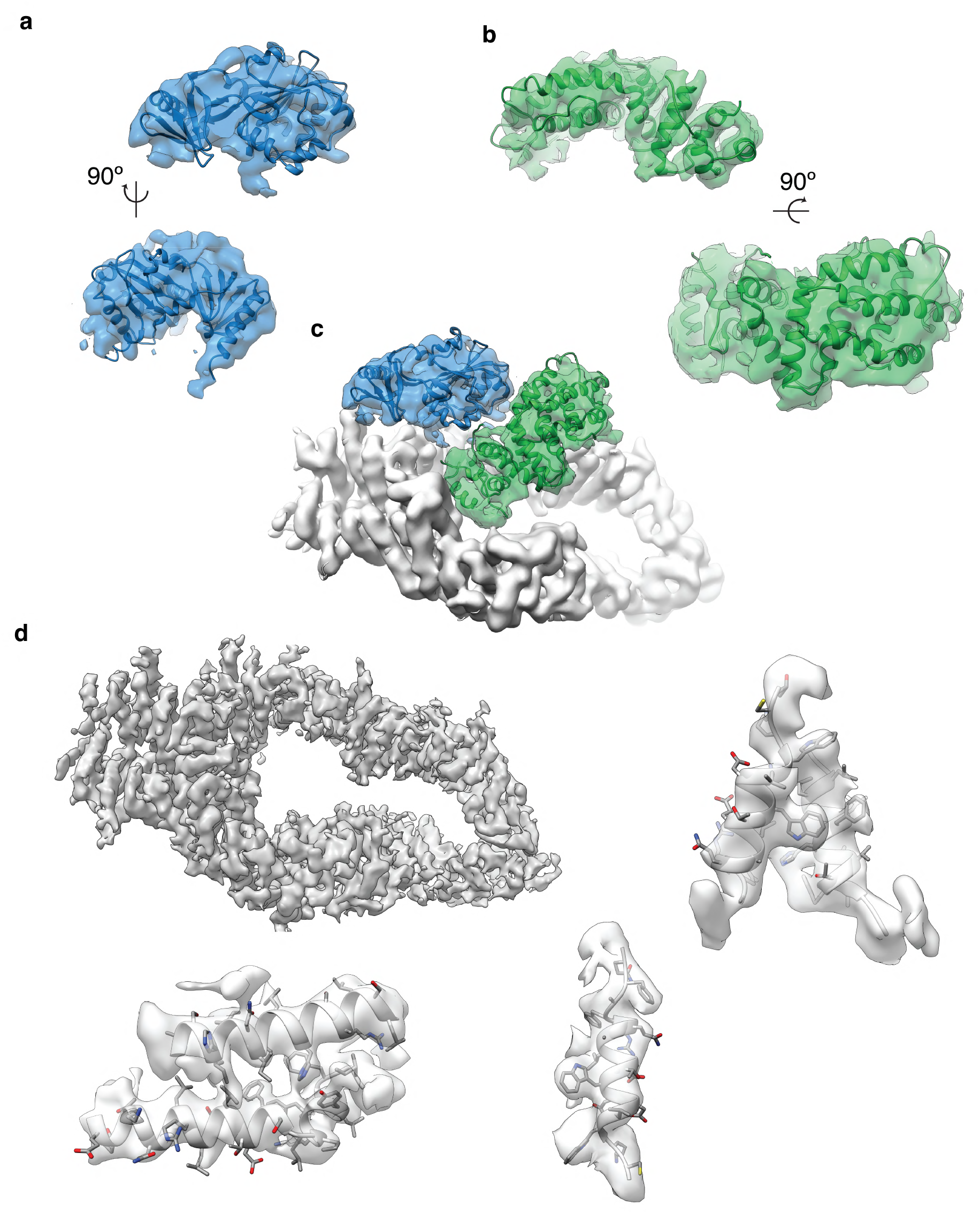
Map to model fit. **a.** Map-to-model fit of the SEC-PH domain (blue). **b.** Map-to-model fit of the GRD domain, clearly resolved secondary structure is evident. **c.** The autoinhibited conformation of NF1, GRD (green) and SEC-PH (blue) domains are well resolved against the scaffold. **d.** The sharpened 4.6 Å reconstruction of the core NF1 scaffold and select regions of density illustrating clearly resolved side chains.

**Fig. S6.**
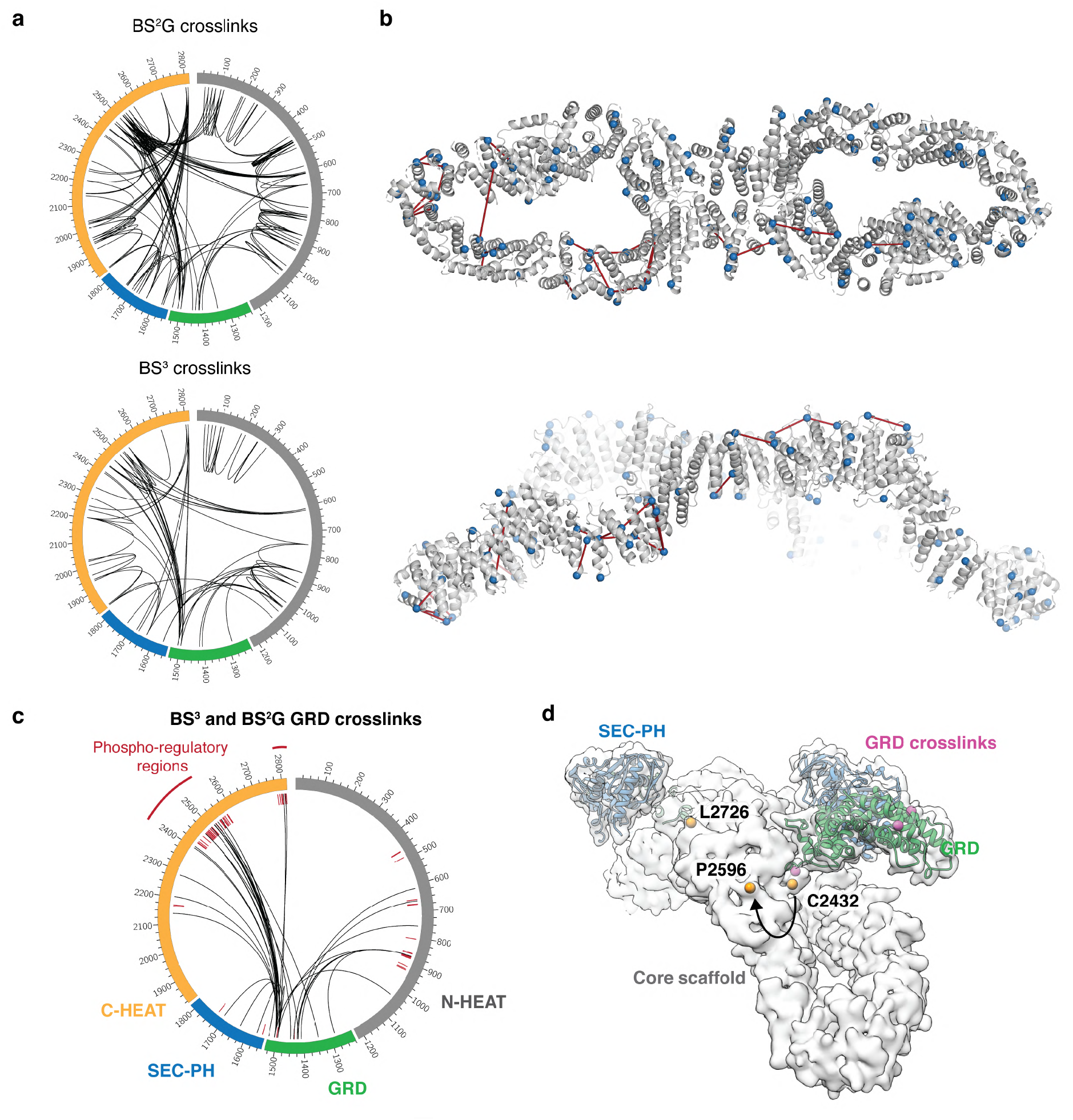
Crosslinking mass spectrometry of NF1. **a.** Crosslinking mass spectrometry identified a total of 174 BS^2^G crosslinks and 72 BS^3^ crosslinks within the NF1 dimer. **b.** Mapping of the BS^2^G crosslinks onto the NF1 structure revealed that the majority of all crosslinks mapped to allowable regions of the NF1 model. C**α** carbons of crosslinked residues are shown as spheres and lines indicate crosslinked residues. **c.** Circos plot highlighting all BS^3^ and BS^2^G crosslinks observed between the GRD domain and the rest of the NF1 molecule. GRD crosslinked regions map remarkably well to the two known NF1 phosphorylation clusters(51). Known NF1 phosphorylation sites(52) are indicated as dashed red lines on the inside of the circos plot. **d.** Although these regions are too flexible to appear in the cryo-EM maps, they are each situated adjacent to the GRDs and appear poised to further regulate either the conformation of the GRD or membrane binding capacity. Structural representation of the NF1 surface with the GRD and SEC-PH domains in cartoon format. The C**α** carbon of the exit (C2432) and entry (P2596) residue of the main phospho-regulatory loop are indicated as yellow spheres. The C**α** carbon of the final resolved residue of the C-terminus (L2726) is also indicated as a yellow sphere. C**α** carbons on the GRD that crosslink to the phosphoregulatory loops are indicated as pink spheres.

**Fig. S7.**
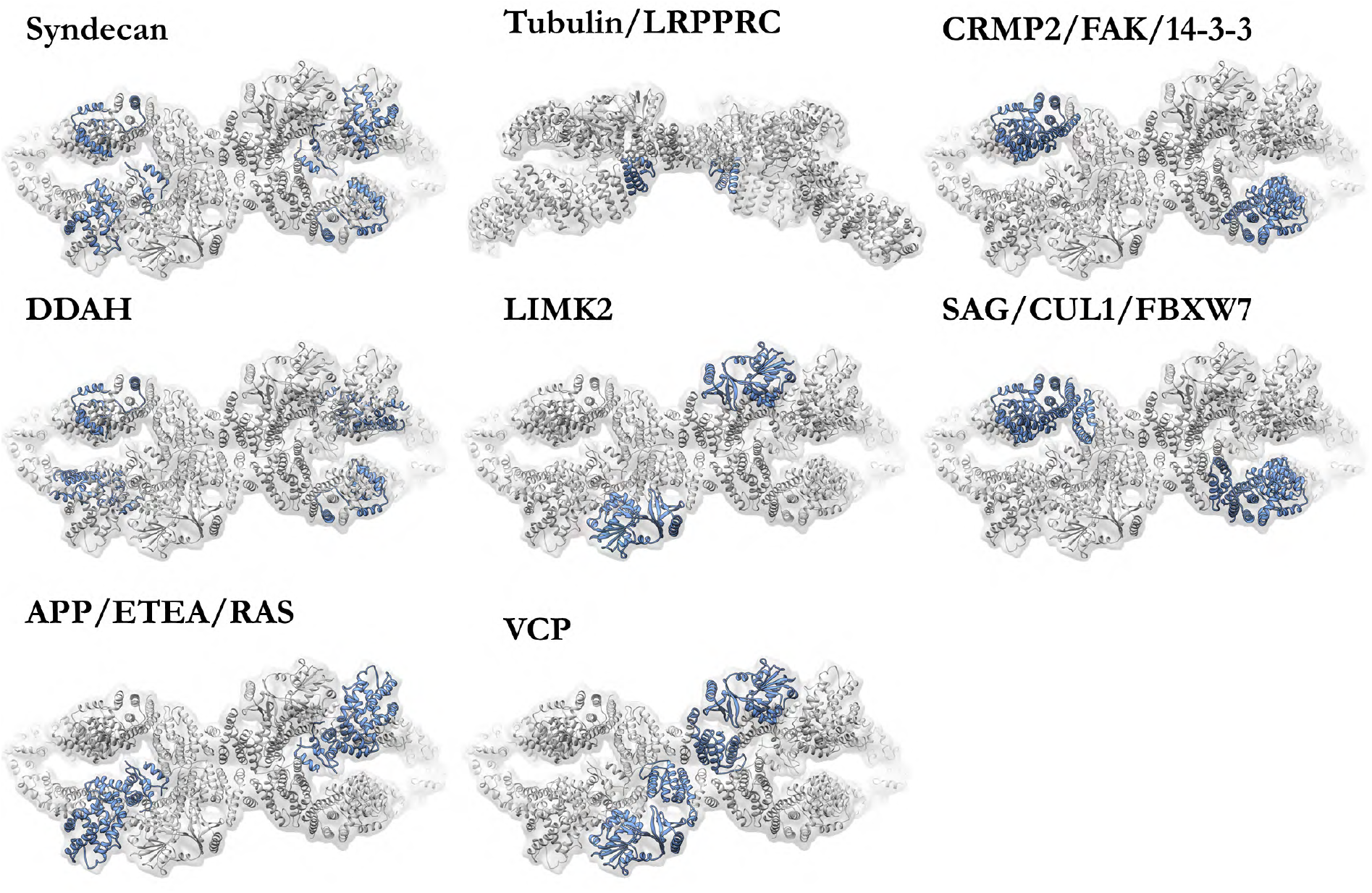
Mapping of known binding sites to the NF1 structure. The position of select binding partners of NF1 (grey) are mapped onto the structure (blue)(2).

**Fig. S8.**
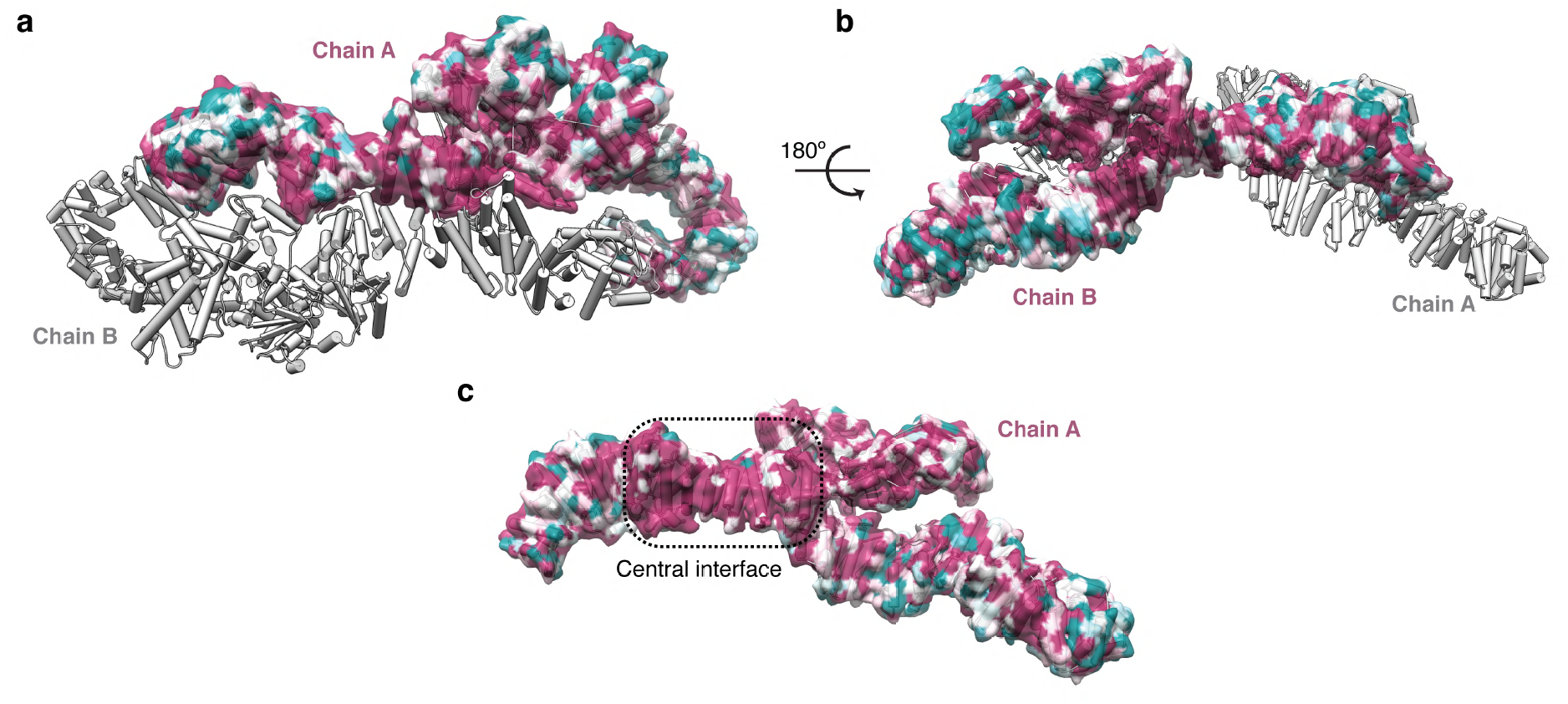
Surface conservation of NF1. NF1 surface conservation analysed using ConSurf(53). **a.** The NF1 dimer with surface conservation mapped to a single chain. **b.** NF1 rotated 180° to show surface conservation of the second chain. **c.** A single NF1 chain displayed to show the conserved central interface.

**Fig. S9.**
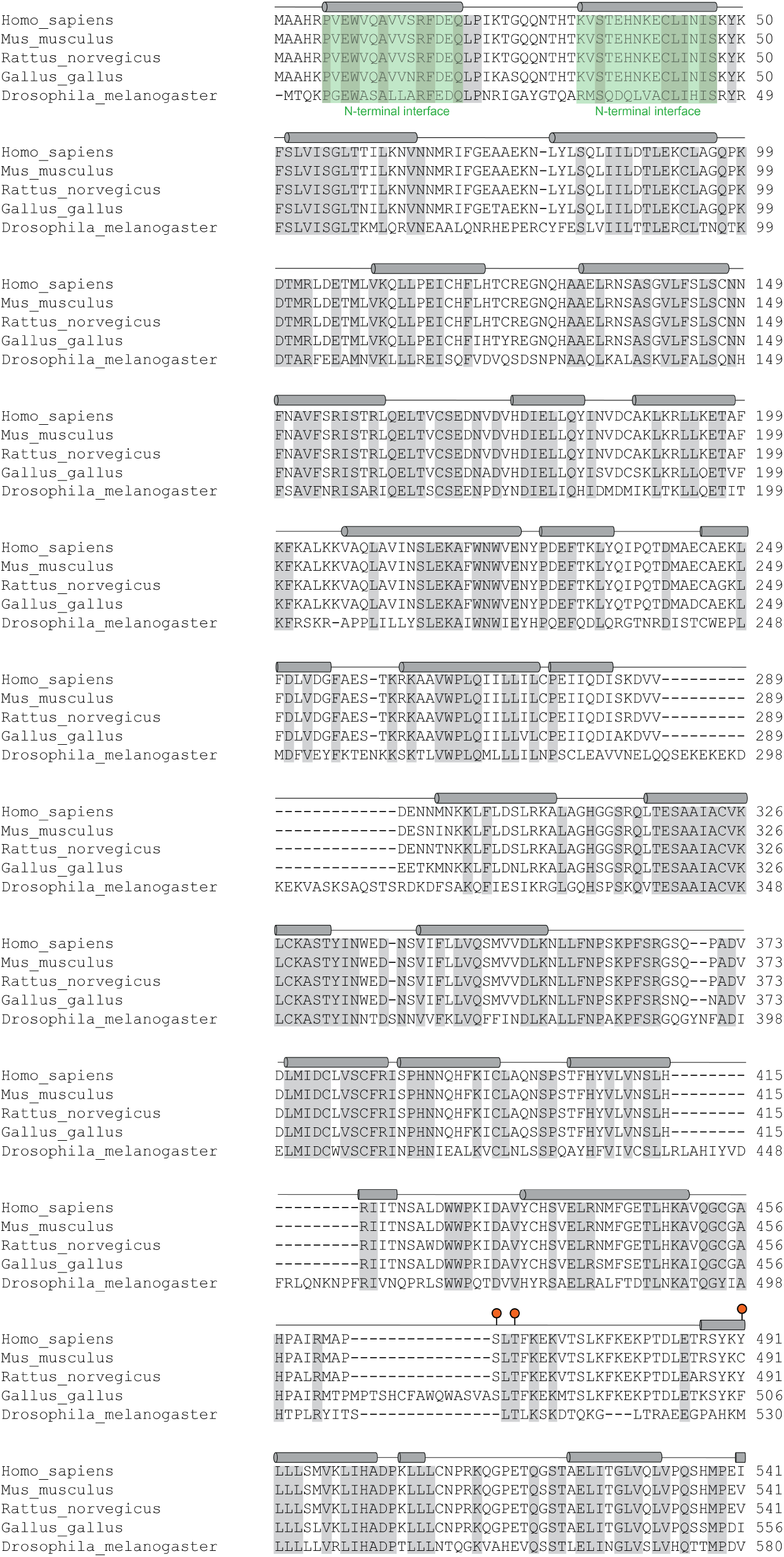
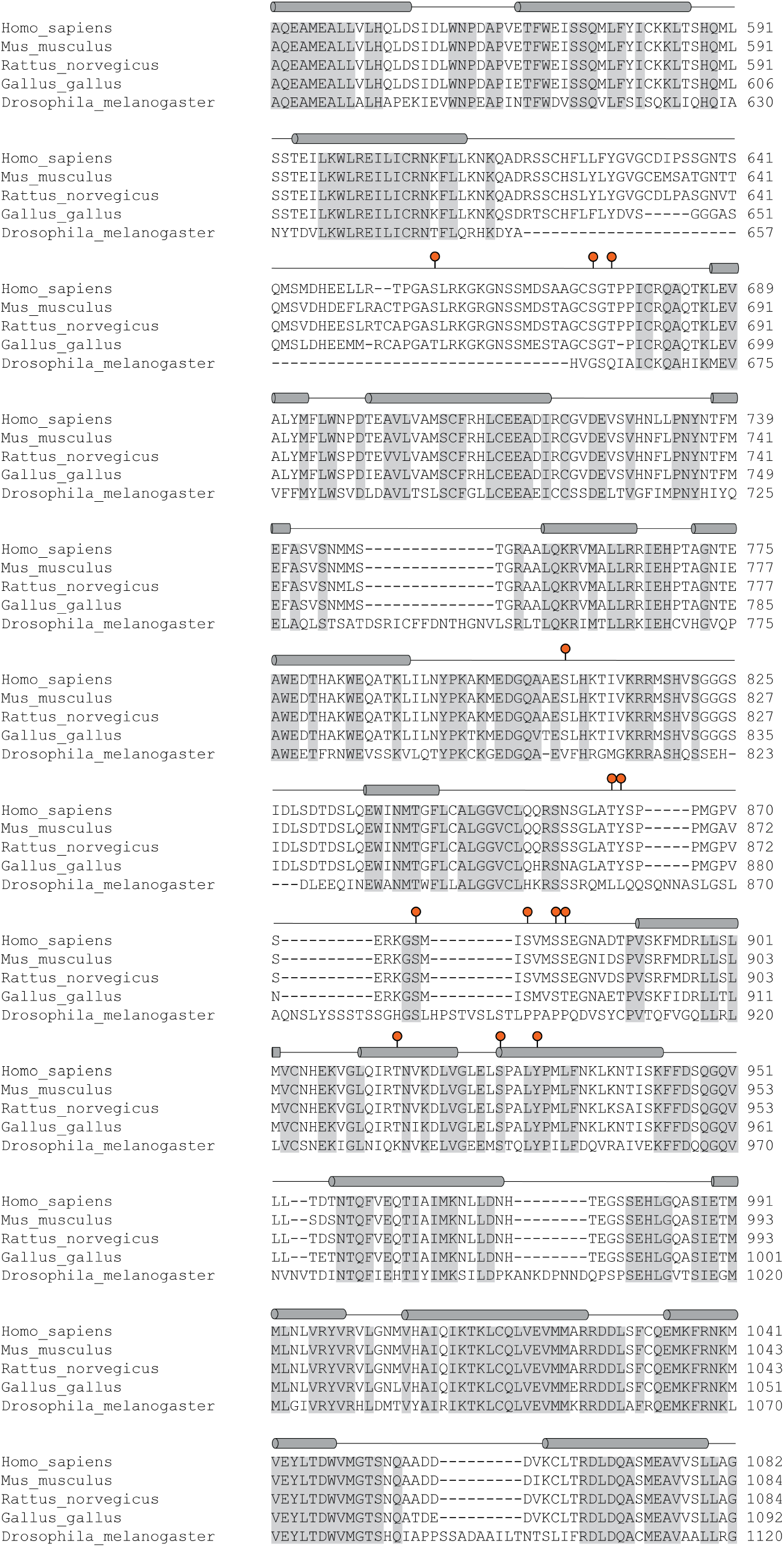
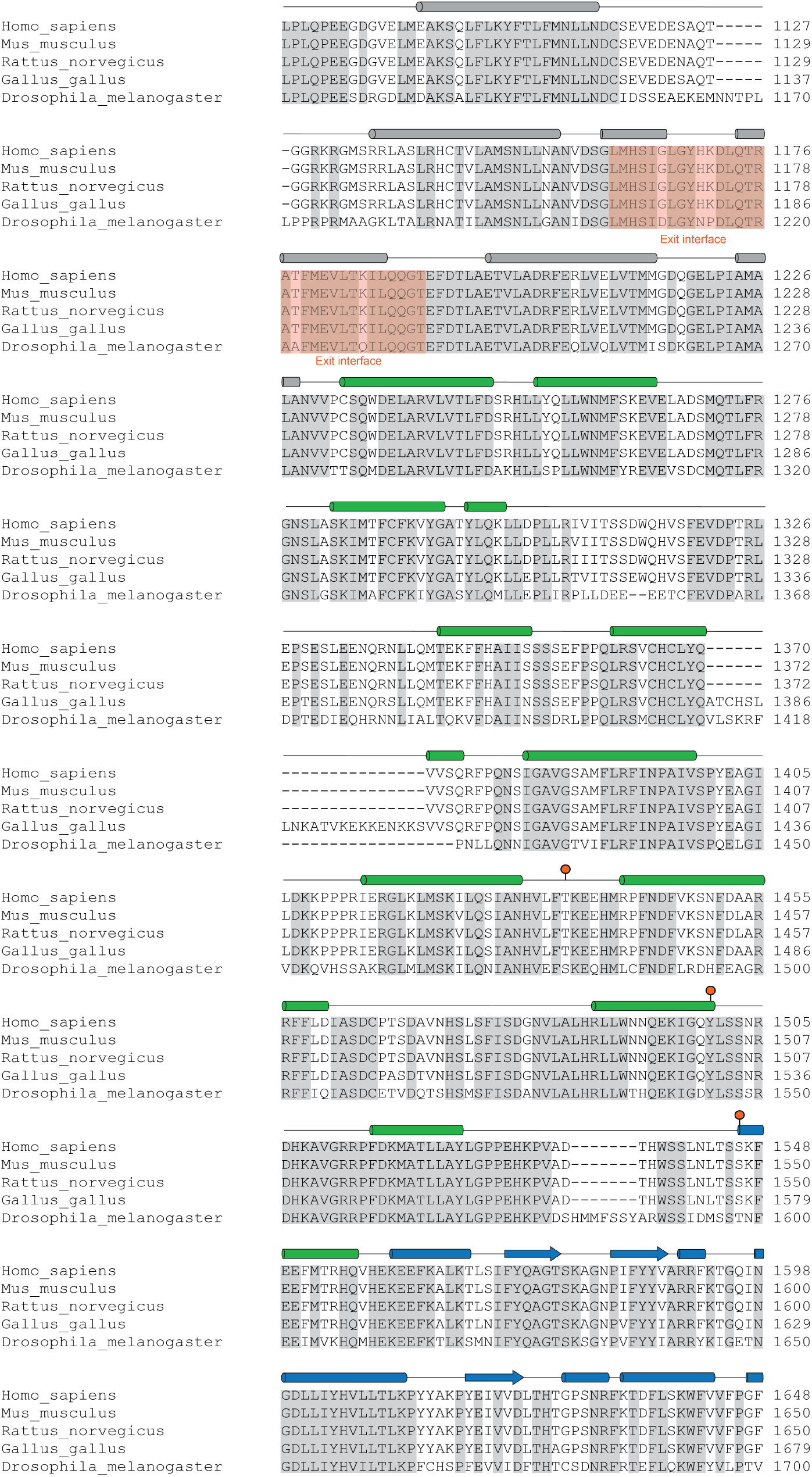
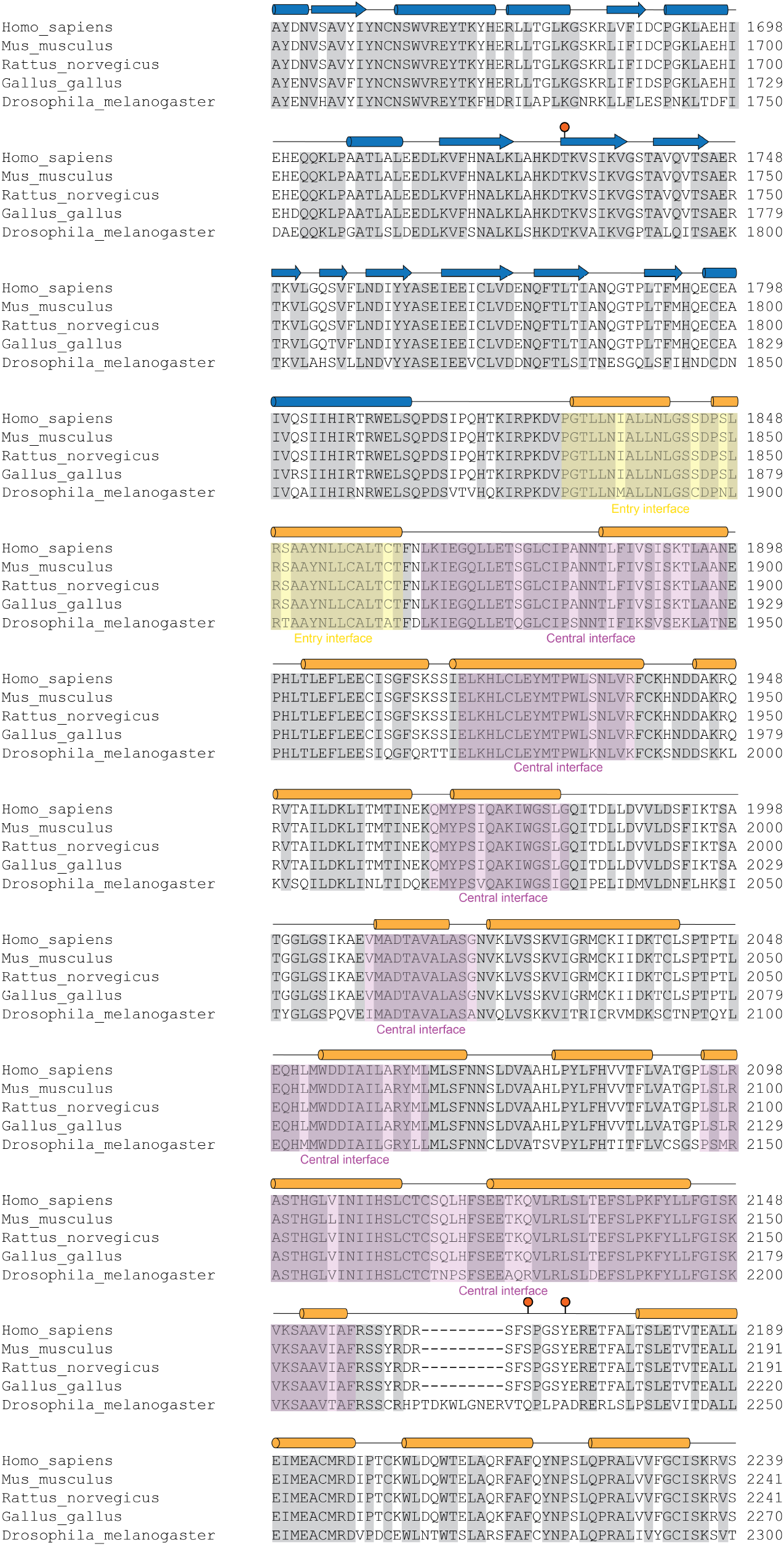
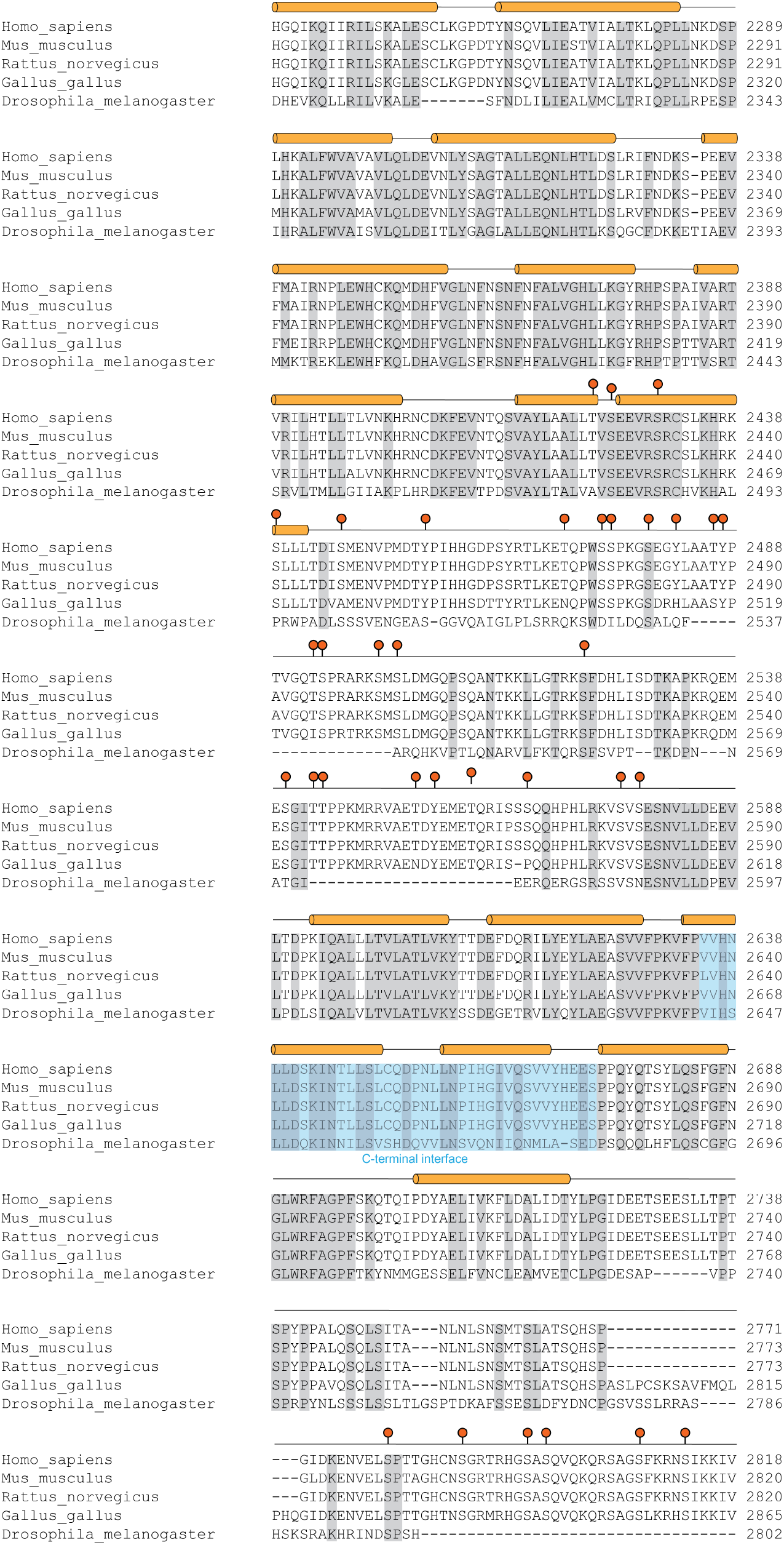
NF1 multiple sequence alignment and secondary structure assignment. Multiple sequence alignment of NF1. Conserved regions are highlighted in grey with the secondary assignment from the NF1 structure indicate above the alignments. Key sites of phosphorylation(51) and dimer interfaces are indicated.

**Table S1.**
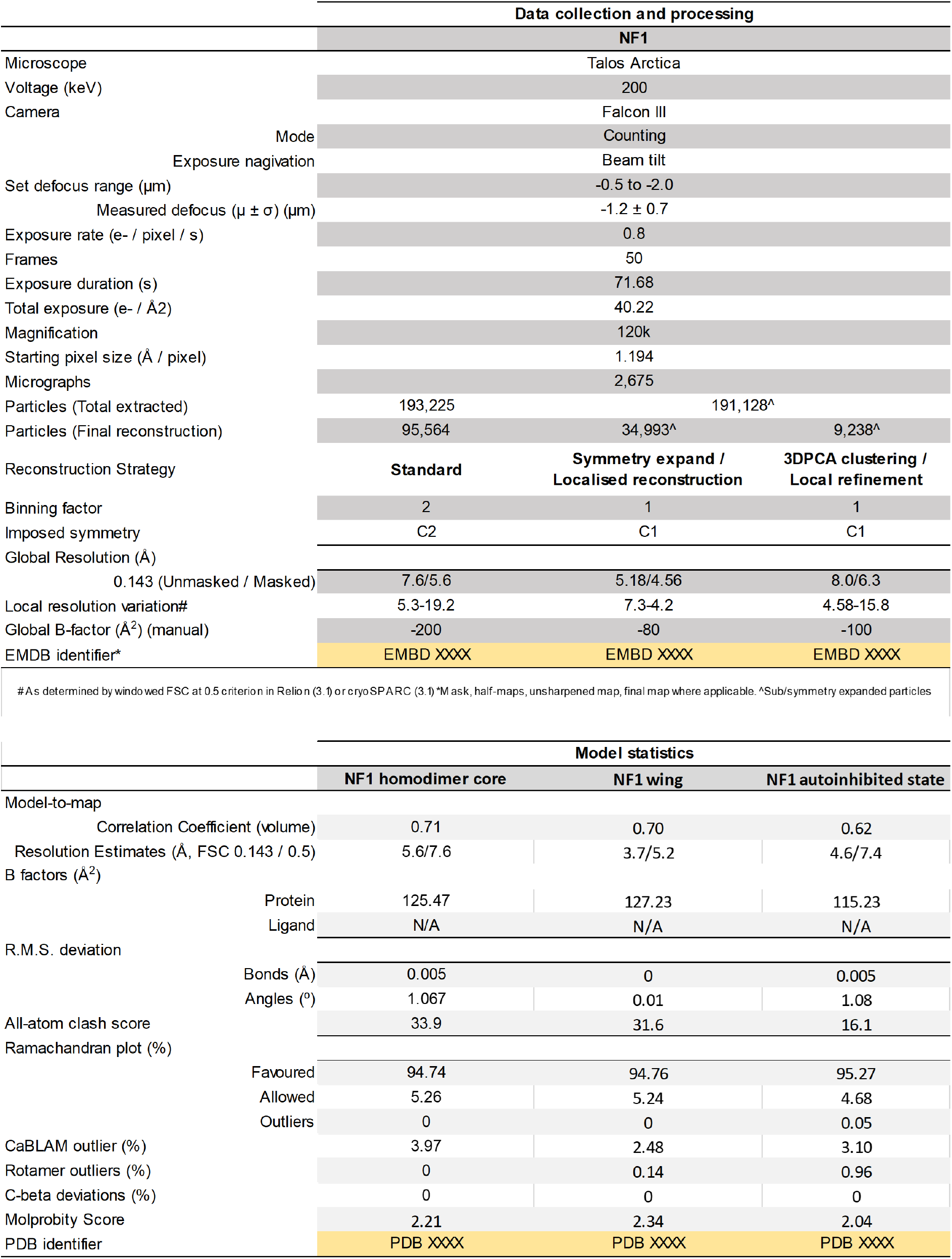
Cryo-electron microscopy data collection, processing and analysis statistics.

**Table S2.**
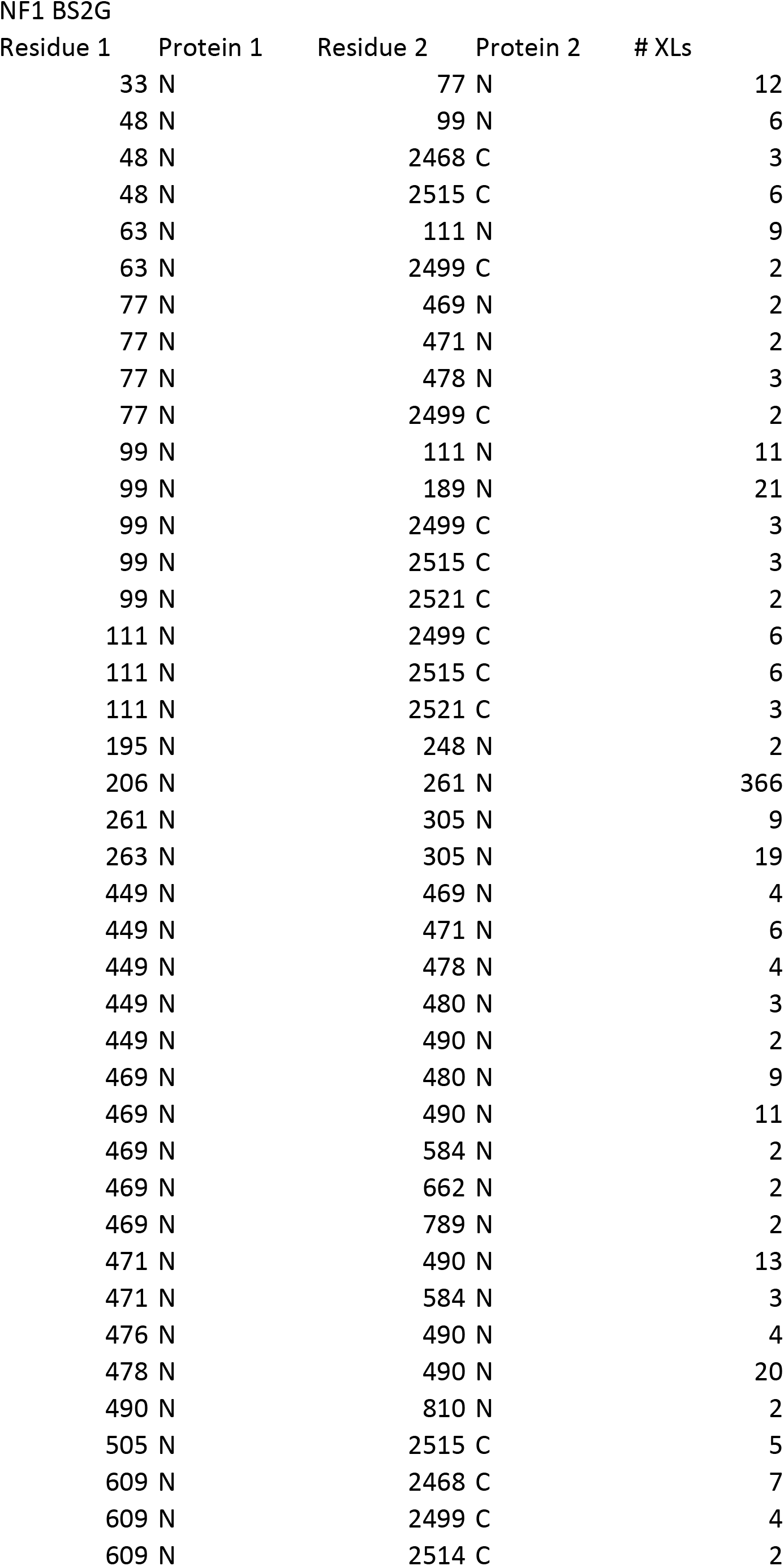
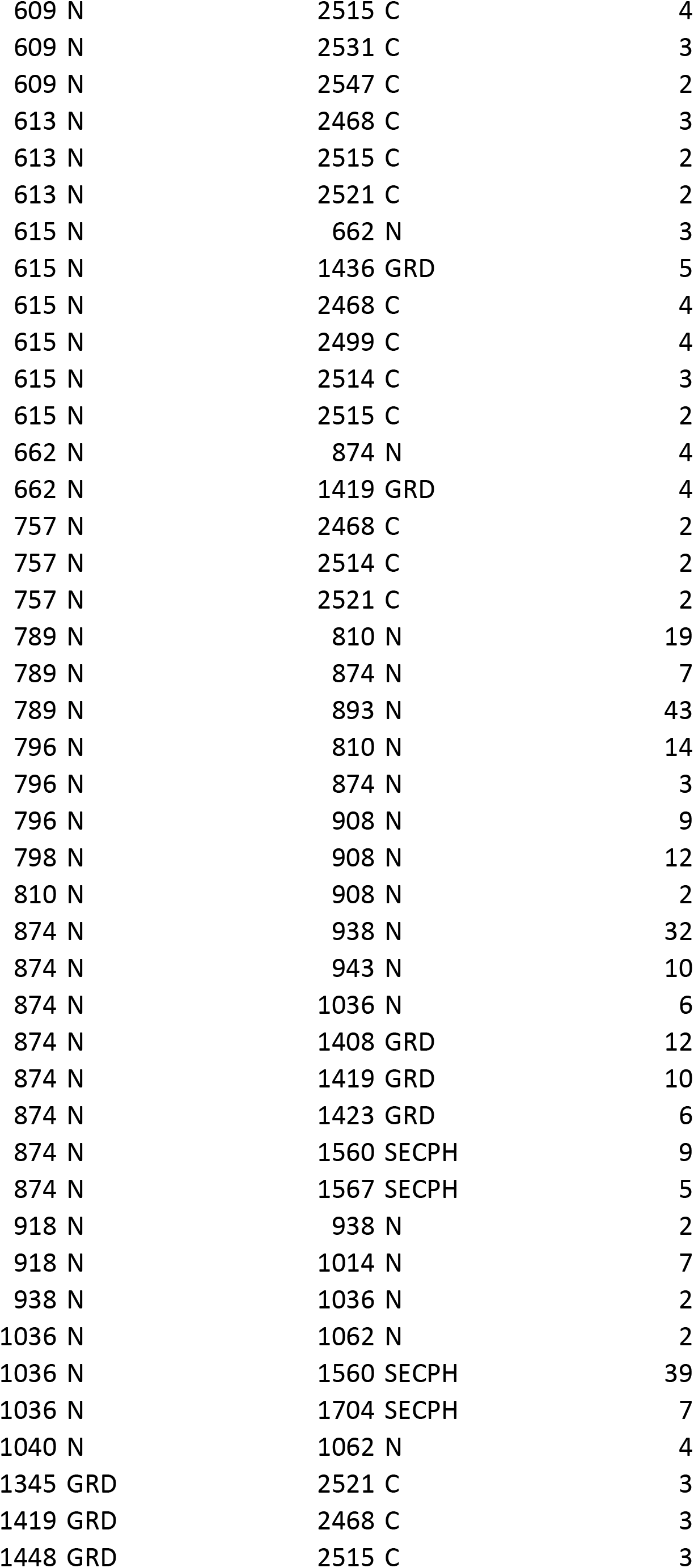
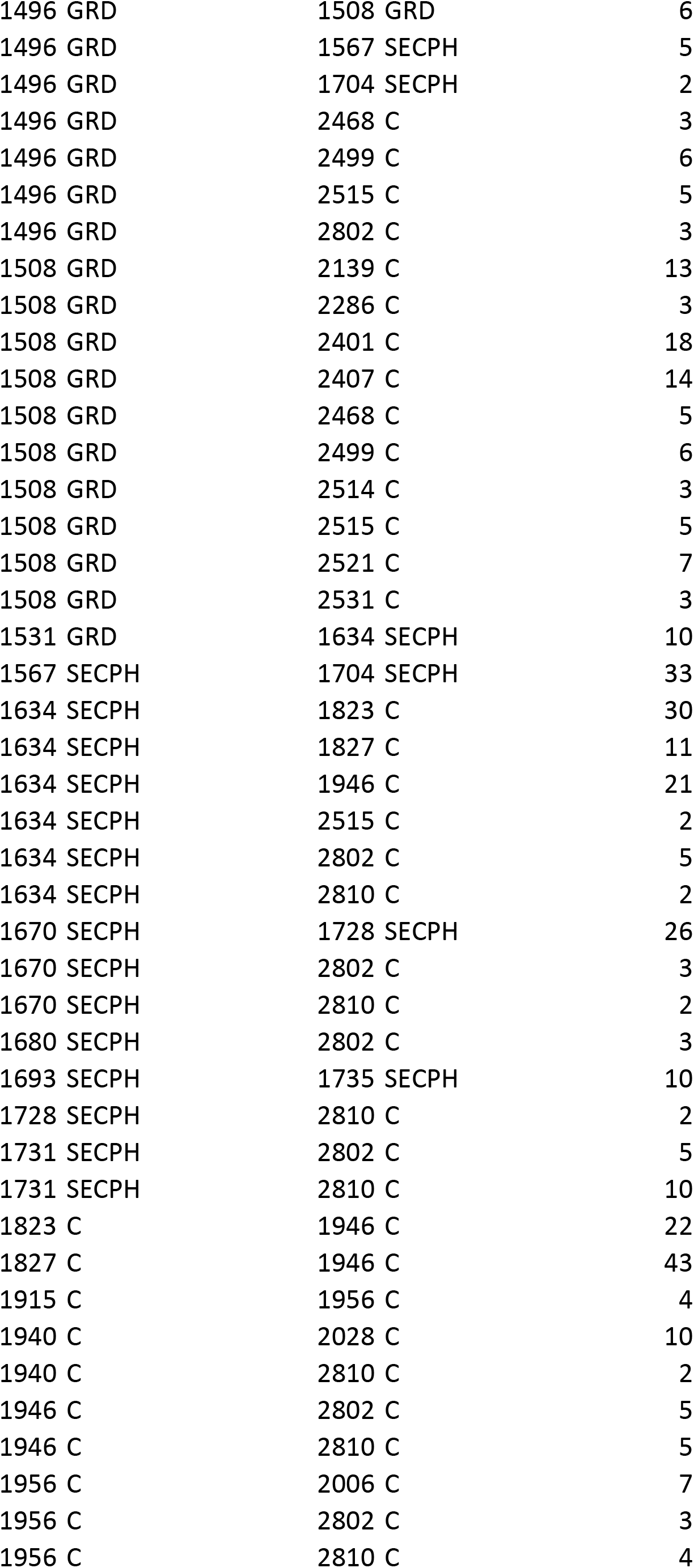
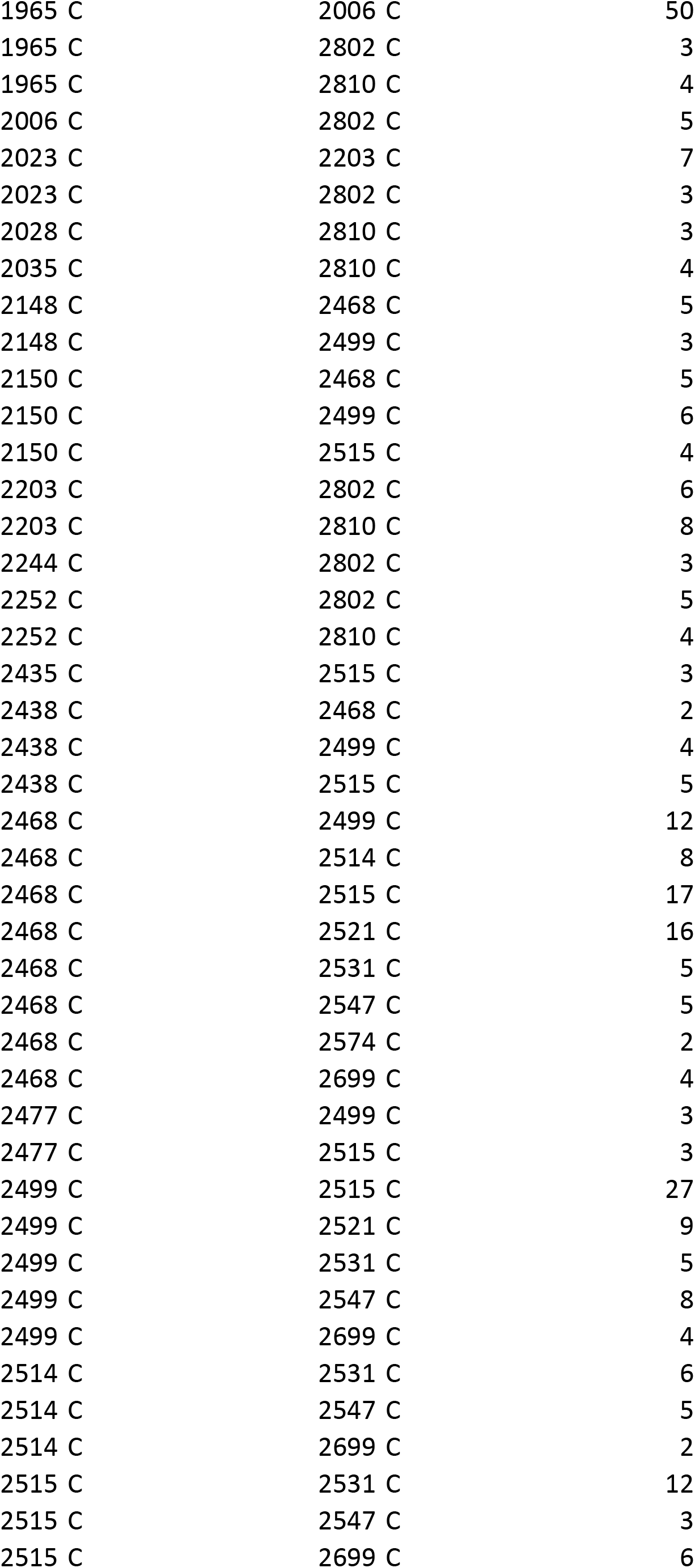
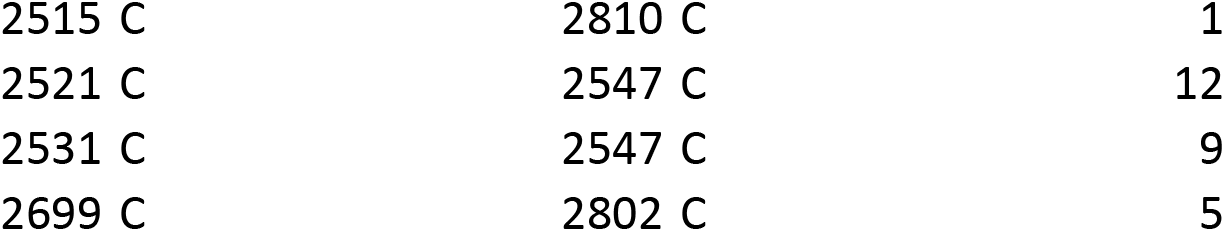
NF1 crosslinking mass spectrometry sites. **See Extended_data_table_2.xlsx**

## Notes

### Competing Interest Statement

The authors have declared no competing interest.

